# A scalable *in vitro* cell competition model identifies preleukemic super competitor clones

**DOI:** 10.64898/2026.10.21.760611

**Authors:** Lilia Cabrera-Cosme, Juan A. Rubio-Lara, Ellie Bennett, Joanna Miłek, Chloë Baldreki, Adam Dowle, M.S. Vijayabaskar, Sophie J. Davies, Ewan Taylor-Smith, Fiona M. Bain, Sandy MacDonald, Sally James, Karen Hogg, Maria Jassinskaja, Berthold Gottgens, Nicola K. Wilson, David G. Kent

## Abstract

Preleukemic and leukemic conditions are frequently characterized by the acquisition of recurrent genetic mutations in hematopoietic stem cells (HSCs). One of the most frequently mutated genes in both clonal hematopoiesis and myeloid malignancies is the ten-eleven translocation 2 (TET2) gene, which encodes an epigenetic enzyme with a critical role in the regulation of gene transcription. While substantial work has shown that TET2 loss-of-function mutations lead to a clonal advantage, the underpinning cellular mechanism remains unclear, especially with respect to the early interactions with other cells. To study these early cell interactions of mutant and non-mutant cells, we developed an *in vitro* clonal competition platform where single-cell derived cultures are split into subclonal fractions which can be simultaneously cross-competed against multiple different clones of an opposing genotype. We identify a novel preleukemic cell phenotype that parallels the developmental biology phenomenon of super-competition, whereby ∼10% of TET2-mutant clones actively and selectively suppress wild type (WT) cell expansion and, in some cases, completely eliminate WT clones. Molecular profiling and functional assays point to RAC1-GEF signalling inhibition as a specific WT cell vulnerability that is promoted, and tolerated, by neighbouring TET2-mutated clones. In addition to pinpointing selective drivers of non-mutant cell suppression, this clonal competition platform offers a broadly applicable new approach to study cell competition dynamics at an unprecedented scale.

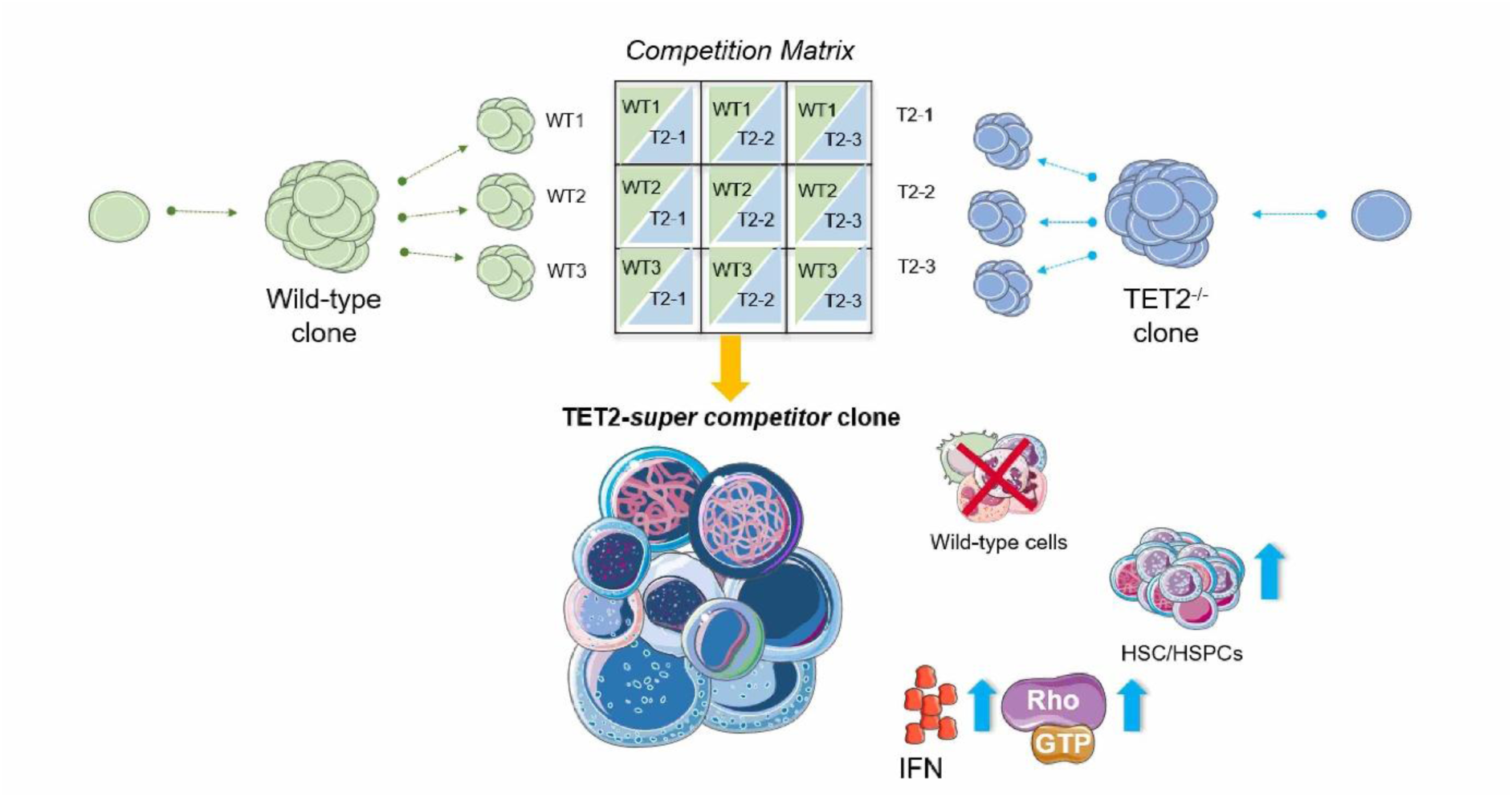

*Key points:*

- Split clonal culture system allows for the modelling of clonal competition *in vitro* between a single clone against multiple clones of an opposing genotype
- A subset of TET2-mutant HSCs are “super competitors”, actively suppressing wild-type cells.
- TET2-super competitors increase phenotypic HSCs and tolerate Rac1-GEF signalling disruption.

## Introduction

Blood cancers have long been viewed as the products of cellular genetic evolution, with natural selection underpinning the outgrowth of the leukemic clone (Nowell, 1976; Landau et al, 2014; Ferrando & Lopez-Otin, 2017; Lyne et al, 2021). Recurrent genetic driver mutations have been identified, conferring on hematopoietic stem cells (HSCs) a competitive advantage over their non-mutant counterparts (Watson et al, 2020; Machado et al, 2023; Papaemmanuil et al, 2013; Papaemmanuil et al, 2016). Building upon this, clonal hematopoiesis (CH) was also first described as a clonal gain-of-fitness with increased risk of hematological malignancies and adverse clinical outcomes (Jaiswal et al, 2014; McKerrel et al, 2015; Bowman et al, 2018). Among the mutated genes driving competitive advantage and enhanced self-renewal is TET2, an epigenetic regulator commonly mutated in both CH and a wide range of myeloid malignancies (Delhommeau et al, 2009; Ortmann et al, 2015).

Clonal competition is typically driven either by an intrinsic proliferation advantage (e.g., clone grows faster or survives longer), or an extrinsic supportive stimulus (e.g., a specific growth factor) which leads to clonal outgrowth and/or survival (de Haan & van Zant, 1997; Zon et al, 2008; Li, 2011; Lee et al, 2017; Lee-Six & Kent, 2020; Kent, 2022). In the developing organism and during tumorigenesis, however, a common mechanism of clonal competition is the active suppression of neighbouring cells (e.g., successful competition achieved through a relative, rather than an absolute, clonal advantage). This question is not experimentally tractable at the level of single cells in the hematopoietic system though, and researchers have therefore left this cellular mechanism relatively underexplored. With growing evidence that systemic environmental triggers such as inflammation and radiation can shift the fitness of individual hematopoietic clones, this warrants further investigation (Bondar & Medzhitov, 2010; Marusyk et al, 2010; de la Cova et al, 2014; Fabre et al, 2022; Chen et al, 2026). In particular, both mouse and human TET2-mutated hematopoietic stem and progenitor cells (HSPCs) promote inflammatory stress via IL-1β (McClatchy et al, 2023; Huerga-Encabo et al, 2026) and TET2-mutated human acute myeloid leukemia (AML) cell lines have a chemotherapy-response advantage (Morinishi et al, 2020). This further accords with mouse models where TET2-deficient cells gain a competitive advantage through increased MPL signalling (Yang et al, 2025) and increased inflammatory markers are observed at the transcriptomic and proteomic levels (Jassinskaja et al, 2026). What remains unclear though, is whether TET2-deficient cells outcompete non-mutant cells solely via elevated proliferation/survival, or if they actively modulate growth of non-mutant cells. While barcoding and lineage tracing tools exist to measure population dynamics, virtually no tools exist to measure individual clone:clone interactions and the impact on competition dynamics.

To address this question, we adapted the PVA-based HSC expansion system (Wilkinson et al, 2019; Wilkinson et al, 2020, Che et al 2022) to demonstrate that both bulk and single HSC-initiated cultures can be split into molecularly and cellularly equivalent fractions, providing replicate cultures for experimental perturbation. Leveraging this capability, we developed a preleukemic clonal competition assay that placed single TET2-mutated HSC-derived clones in competition against multiple WT clones simultaneously, and vice versa. By tracing the competitive dynamics through different timepoints, we identify a “super-competitor” behaviour in TET2-mutant HSCs, consistently across hundreds of individual competition assays. This behaviour was primarily characterized by the selective depletion of WT cells, despite matched controls showing uninhibited growth. This mechanism of active and selective suppression was explored on a molecular and functional level, identifying disruption of RAC1-GEF signalling as a WT-specific cell vulnerability, which is promoted and tolerated by preleukemic TET2-mutant clones.

## Methods

### Mice

WT and TET2 mutant mice aged 8-12 weeks were used as donors and WT mice (lethal irradiation) and W^41^/W^41^ (sublethal irradiation) were used as transplantation recipients. Strain information and ethical approvals are fully detailed in Supplemental Methods.

### Isolation and expansion of hematopoietic stem cells

ESLAM-Sca-1^high^ HSCs were isolated with experimental details available in the Supplemental Methods. Single-cell or bulk cultures in the PVA expansion system were carried out as previously described (Wilkinson et al, 2020) with additional details available in the Supplemental Methods.

### Splitting of single-cell derived clones

Single-HSC derived clones were split on day 14 across four equivalent fractions into new 96-well, fibronectin coated plates. Full details are provided in the Supplemental Methods.

### Cell competition assays

Following clone splitting, subfractions of WT (CD45.1) or TET2 (CD45.2) clones were mixed at equal proportions to establish clonal competition assays. Full details are provided in Supplemental Methods and in Results.

### Flow cytometry profiling of expanded clones

Day 28 cultures were profiled by flow cytometry to assess total cellularity and cellular composition, with full details in the Supplemental Methods.

### Single-cell RNA-sequencing

Day 28 split fractions for both clonal and bulk cultures were harvested on day 14 and washed with PBS. Libraries were prepared using Evercode WT reagents (Parse Biosciences), according to the manufacturer’s protocol. Further details are included in the Supplemental Methods.

### Bulk RNA-sequencing

Day 28 control wells of competition cultures were harvested and sequenced for RNA-sequencing with full details in the Supplemental Methods.

### Data-Independent Acquisition Mass Spectrometry (DIA-MS)

Supernatants of non-competed control wells were harvested on days 26 and 28 to assess the secretome. Full details are included in the Supplemental Methods.

### HSC transplantation

*In vivo* HSC transplantation assays were undertaken for both control split cultures and for control wells from competition cultures on day 28. Full details are in the Supplemental Methods.

### *In vitro* validation of targets

Bulk HSC cultures containing 25 cells per well were treated with Rho-GTPase inhibitors or recombinant proteins in order to validate the findings from the transcriptomic and secretomic profiling of competitor clones. Full details are in the Supplemental Methods.

## Results

### Split clones are phenotypically, functionally and molecularly equivalent

PVA-based mouse HSC expansion cultures (Wilkinson et al, 2019; Wilkinson et al, 2020) exhibit substantial heterogeneity at the single cell level, which modified *in vitro* HSC isolation strategies only partially resolve (Che et al, 2022; Becker et al, 2023; Zhang et al, 2024), leaving input HSC heterogeneity as a substantial obstacle. Comparative experiments that require experimental perturbations to start from genuinely equivalent populations are therefore challenging. To address this, we established an approach where a single HSC-derived clone is physically divided into four sister fractions which could each be simultaneously exposed to different downstream conditions, while also linking back to the same parental HSC clone. In order for this approach to be robust, sister wells must be roughly equivalent at the population level.

To establish split culture equivalence, single long-term HSCs (ESLAM HSCs-CD45^+^EPCR^+^CD150^+^CD48^-^) (Kent et al, 2009) were expanded for 14 days, after which they were harvested and split evenly across four new sister wells (25% of original clone per well) and expanded for a further 14 days (**Figure 1A**). Flow cytometric profiling showed consistent number and proportions of *in vitro* expanded phenotypic HSCs, as measured by EPCR+LSK (ELSK) frequencies and counts (Che et al, 2022), as well as equivalent numbers of viable cells across sister wells (**Figure 1B**). This was also demonstrated for TET2 sister wells, wherein ELSK frequencies were equivalent across split clone fractions (**Supplementary Figure 1A**). The intraclass correlation coefficient (ICC) of ELSK frequencies was 0.90, demonstrating strong consistency across split fractions of 38 parent HSC clones (Koo & Li, 2016) (**Figure 1B**). ICC for ELSK counts across split fractions was 0.80, and 0.89 for viable cell counts, which further validated the consistent phenotypic composition of sister wells. A one-way analysis of variance (ANOVA), showed that 89.88%, 80.03% and 89.13% of data variation in ELSK frequencies, counts and viable cell counts -respectively - came from between cultures rather than within individual split clones (**Supplementary Figure 1B**). The proportions of Gr-1^-^ CD11b^-^ myeloid cell lineage-negative (Lin-) and Lin-Sca-1^+^c-Kit^+^ (LSK) cells were also consistent across sister wells (**Figure 1C**), a pattern also observed in TET2-mutated single-HSC split clones (**Supplementary Figure 1B**). We also measured long-term reconstitution potential across 12 sister wells from 3 clones, finding equivalent donor cell repopulation across sister wells (**Figure 1D**).

**Figure 1.**
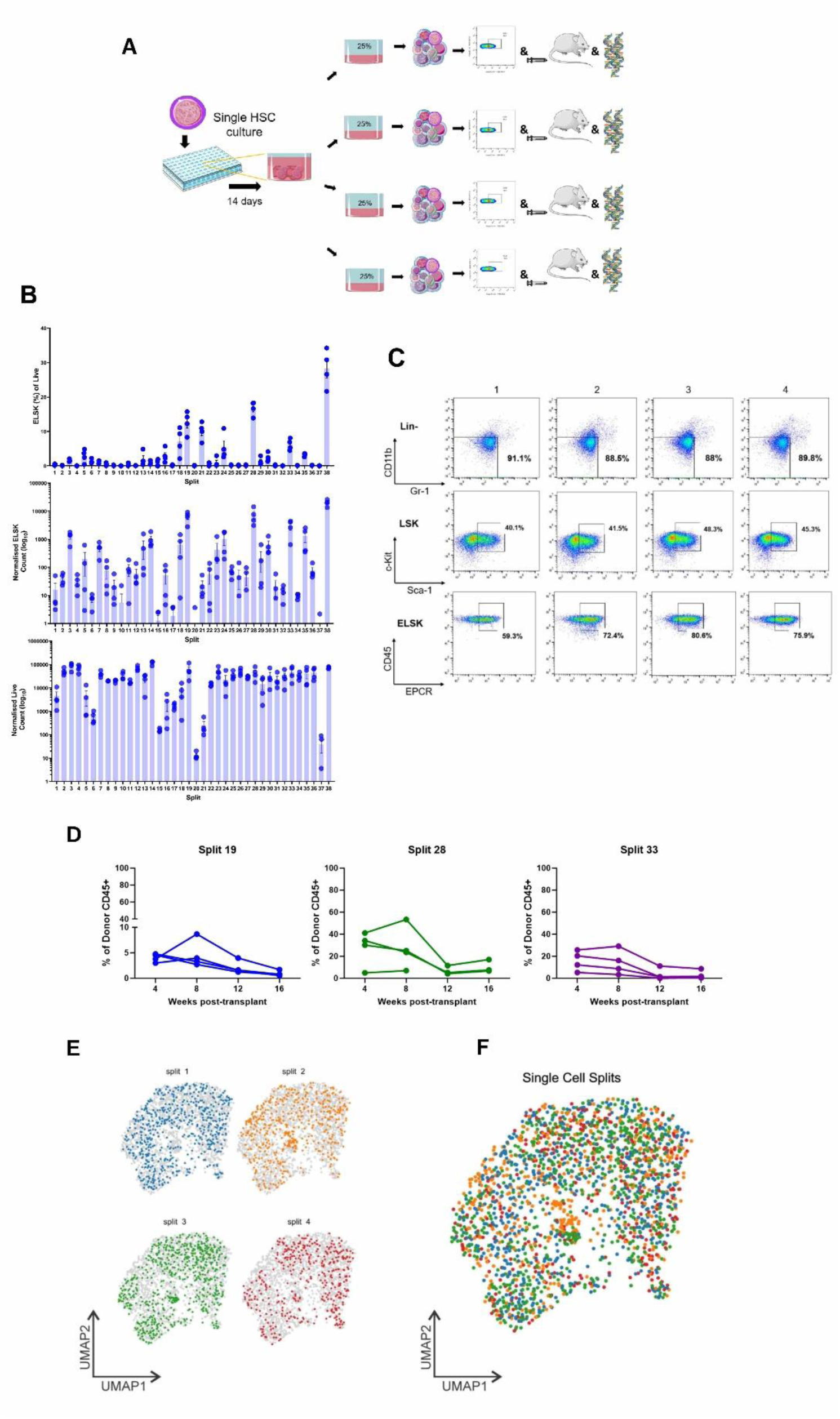
Single-cell split cultures are phenotypically, functionally and molecular homogeneous. **A)** Experimental design for split cultures. Single HSCs were isolated from WT mice (n = 3) and expanded for 14 days and then fractioned into four equal portions across four wells of a new plate. Resulting sister wells were cultured for 14 days (28 days total) and harvested for flow cytometry, transplantations and scRNA-seq. **B)** Frequencies of phenotypic HSCs (defined as ELSK cells) (top), counts (middle), and viable cell counts (bottom) in sister well (n=38). Each number represents a set of four sister wells derived from the same original clone. Mean and standard deviation between ELSK frequencies, counts and viable cell counts within split cultures were similar across sister wells. Bars represent SEM. **C)** Sister well phenotypic analysis for group #38 shown in **B**. Lin^-^(top) defined as Gr-1^-^CD11b^-^. LSK and ELSK (middle and bottom, respectively). **D)** *In vivo* functional validation of split culture homogeneity where donor chimerism (%) in the CD45^+^ population for three split-culture recipients are shown. Notably, sister wells showed similar repopulation kinetics throughout the transplantation experiment. There were 4 recipients per well; 20% of total well contents were used for transplantation, and 20% for flow cytometry. Recipient #4 in group #28 died after the 8-week timepoint due to reasons unrelated to the transplant. **E)** Transcriptomic landscape of split cultures corresponding to individual sister wells from clone #38 in **B**. Gene expression was highly similar across split cultures with 60% of total well contents profiled. **F)** Merged UMAP plots displaying the gene expression profiles of all split cultures within the group. Expression of representative genes (**Supplementary** Figure 1C) and clustering (**Supplementary** Figure 1D), were homogeneous across sister wells.

We next compared the molecular composition of sister wells by performing single cell RNA-sequencing on all four sister wells of the clone with the highest ELSK frequency (clone number 38 in Figure 1B) (**Figure 1E-F**). Sister wells were molecularly equivalent, with cells similarly distributed across the transcriptional landscape. We next evaluated expression levels of genes commonly detected in *ex vivo-*expanded HSCs (**Supplementary Figure 1C**) where, again, sister wells had equivalent expression associated with HSC functionality and myeloid differentiation, such as *Mecom* (Voit et al, 2023), *Mpo* (Nauseef et al, 1988), and *Kit*. Bulk cultures could also be split into equivalent cultures using the same analytical approach (**Supplementary Figure 1D**). Collectively, these analyses demonstrate that both single HSC-derived and bulk HSC cultures can be split into equivalent fractions at day 14 and sister wells continue to behave as matched replicates, providing a validated foundational framework for comparative competition experiments.

### TET2 clones can exhibit a super-competitor phenotype characterized by active elimination of WT cells

To directly evaluate competitive behaviour whilst controlling for inter-clone variability, we designed a paired clonal competition assay using ESLAM-Sca-1^high^ HSCs (Wilson et al, 2015) isolated from WT (CD45.1) or TET2^-/-^(CD45.2) (Ko et al, 2011; Shepherd et al, 2018) mice. Single HSCs were expanded for 14 days and split into four sister wells (**Figure 2A**). We designed a competition matrix whereby one sister well from each HSC-initiated culture was kept unmanipulated as a control well and the remaining three sister wells were co-cultured with three sister wells from the opposing genotype at the same time. This allowed for the profiling of competition dynamics in one single clone against multiple different clones, thereby increasing the applicability and statistical power of the assay.

**Figure 2.**
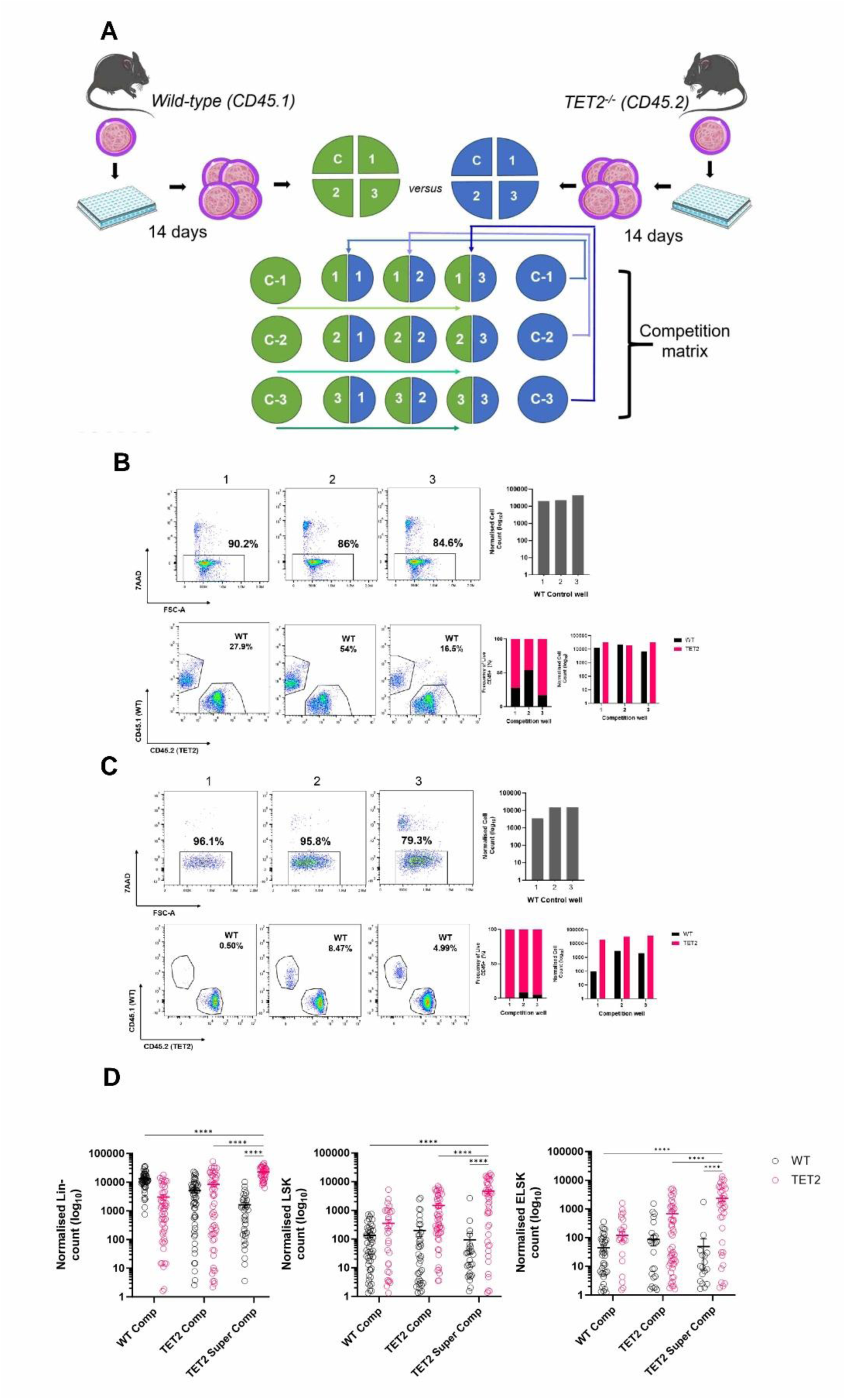
Design of a clonal competition model and the identification of TET2-super competitors. **A)** Experimental design of an in vitro model of cell competition. Single HSCs from wild-type and TET2^-/-^ mice were sorted and expanded independently for 14 days. Clones from each genotype were visually assessed and split four ways. Resulting sister wells were evenly distributed across a competition matrix, a 3x3 arrangement consisting of three control sister wells from each corresponding genotype on each side of the matrix. At the centre of the matrix, sister wells from WT are mixed with sister wells of TET2^-/-^ at a 1:1 ratio (50% of WT and 50% of TET2 by volume). Plates were cultured for a total of 28 days and cultures were then harvested for flow cytometry analysis. **B)** Top, Live cell frequency and cell number in WT-competitor control sister wells (non-competed). Plots are shown for each control (left), together with normalized cell counts (right). Bottom, Example of a TET2-competitor clone. The congenic CD45.1/CD45.2 system allows for the discrimination of WT (CD45.1) or TET2 (CD45.2) cells. Here, flow cytometry plots (left) show a TET2-competitor clone which has outcompeted WT sister wells in two out of three cases. **C)** Top, non-competed WT control sister well whose paired sister wells were outcompeted by the TET2-super competitor clone shown below. Bottom, A TET2-super competitor clone which outcompeted its rivals by a 70:30 proportion across all competition wells within the matrix. Comparison to WT cell number within the competed sister well and its paired control sister well revealed an approximate 10-fold reduction in WT cell number when co-cultured with a TET2-super competitor. **D)** Lin^-^ (left), LSK (centre) and ELSK (right) absolute numbers across three independent experiments by competitor type. In addition to the ability to outcompete wild-type rivals across a matrix, TET2-super competitor clones had elevated HSC/HSPCs *in vitro*. Two-way ANOVA, **** p < 0.0001. Error bars represent the standard error of the mean (SEM).

Each initial HSC-derived clone (WT or TET2^-/-^) was competed against sister wells originating from three different HSCs of the opposite genotype. If TET2 mutations did nothing to alter HSC growth or clonal dynamics, we would observe a roughly equivalent fraction of “winners” and “losers” in the competition matrix. However, more than twice as many TET2^-/-^ clones (68.67%) preferentially outcompeted WT clones (**Figure 2B-C),** in line with the previously described self-renewal advantage of TET2^-/-^ HSCs (Ko et al, 2011; Moran-Crusio et al, 2011; Shepherd et al, 2018). To understand and characterize this phenomenon more completely, we analyzed 140 competition matrices which comprised 420 WT and 420 TET2 individual HSC-initiated clones over 28 days, totalling 1,260 competition wells across four independent experiments.

Each competition well and its accompanying controls (WT and TET2) were analyzed at day 28 with ratios of WT:TET2 cells compared across each competition matrix. For most competition matrices, the three sister wells derived from an individual HSC were a mix of “winners” and “losers”, which indicated no strong competitive advantage of the initial HSC (e.g., not all equally potent and/or expandable HSCs) (Che et al, 2022; Wilkinson et al, 2019). Nonetheless, an intriguing behaviour emerged for ∼10% of clones (83 of 840) whereby all three competition wells for an individual HSC-initiated clone out-competed the cells from the other genotype. The significant majority of these were TET2^-/-^ clones (∼70%) (**Supplementary Figure 2A-B**) with a subset of these (n=27) outgrowing all three competing wells by a greater than 70:30 ratio (as depicted in **Figure 2C**). Particularly striking were a set of TET2^-/-^ clones which we term “super competitors” where, despite WT sister control wells having normal cell proliferation capacity and starting cell numbers, the TET2-super competitor clone actively suppressed all three WT sister wells when cultured alongside them (**Figure 2C**). This phenomenon is powerfully illustrated across the entirety of the competition matrix, where the ratio of cells produced by WT sister wells in the presence of a TET2-super competitor was >20-fold reduced compared to a standard competition well (**Supplementary Figure 2A**).

### Super-competition wells have elevated stem and progenitor cell numbers *in vitro*

We next sought to determine the total the viable, Lin^-^, progenitor cell (LSK) and HSC (ELSK) fractions (**Figure 2D and Figure S2C-E**). Initial HSCs were classified as one of three “competitor types” based on how their sister wells performed in competition against sister wells of the opposite genotype: i) WT competitors (where WT cells outnumbered TET2 cells); ii) TET2 competitors (where TET2 cells outnumbered WT cells); or iii) TET2-super competitor clones (where all TET2 sister wells both outnumbered (>70:30 ratio) and actively suppressed WT cell growth while also producing large numbers of HSPCs). Interestingly, a higher proportion of immature (Lin^-^) cells was observed in wells of the winning genotype (**Figure 2D** (left)), but when numbers of the more immature LSK and ELSK populations were calculated, TET2^-/-^ cells had more of each irrespective of winning genotype (**Figure 2D)**. This was best exemplified in the ELSK fraction where WT competitors were on average outnumbered by ∼20-fold (TET2 competitors) and ∼80-fold (TET2 super competitors). Complementing this, we also calculated frequencies of the granulocyte/monocyte lineages (**Supplementary Figure 2F-H**). In competition wells, the granulocyte frequency was higher in WT competitors compared to TET2-super competitors (p<0.0001) (**Supplementary Figure 2H**). Collectively, these data show that both TET2 and TET2-super competitors were able to maintain a larger stem/progenitor cell pool throughout the cultures irrespective of the presence of WT cells, whereas WT cells were heavily (and negatively) affected by co-culture with a TET2-super competitor clone.

### Kinetics of acquiring the super competitor phenotype

To better understand competition dynamics over time, we competed clones and sampled each competition well on days 3, 7, 10, and 14 post-competition initiation (**Figure 3**). WT competitor and TET2 competitor clones displayed a gradual day-by-day relative increase that would be consistent with a cell proliferation advantage (**Figures 3A** and **Supplementary Figure 3A-B**). TET2-super competitor clones on the other hand were more heterogeneous in terms of proliferation dynamics with just two clones starting and ending as super competitors (**Figure 3B**) and six of eight clones changing their competition phenotype over time. Four clones that were initially classified as “Tet2 competitor” (**Supplementary Figure 3A)** and two as “non-competitor” phenotypes transitioned into super-competitors by day 14 of the culture (**Figure 3C**). The phenotype transition is most evident when comparing the dynamics of a TET2-super competitor clone which maintained its phenotype (**Figure 3D**) with the one that attained a super competitive advantage across time (**Figure 3E**).

**Figure 3.**
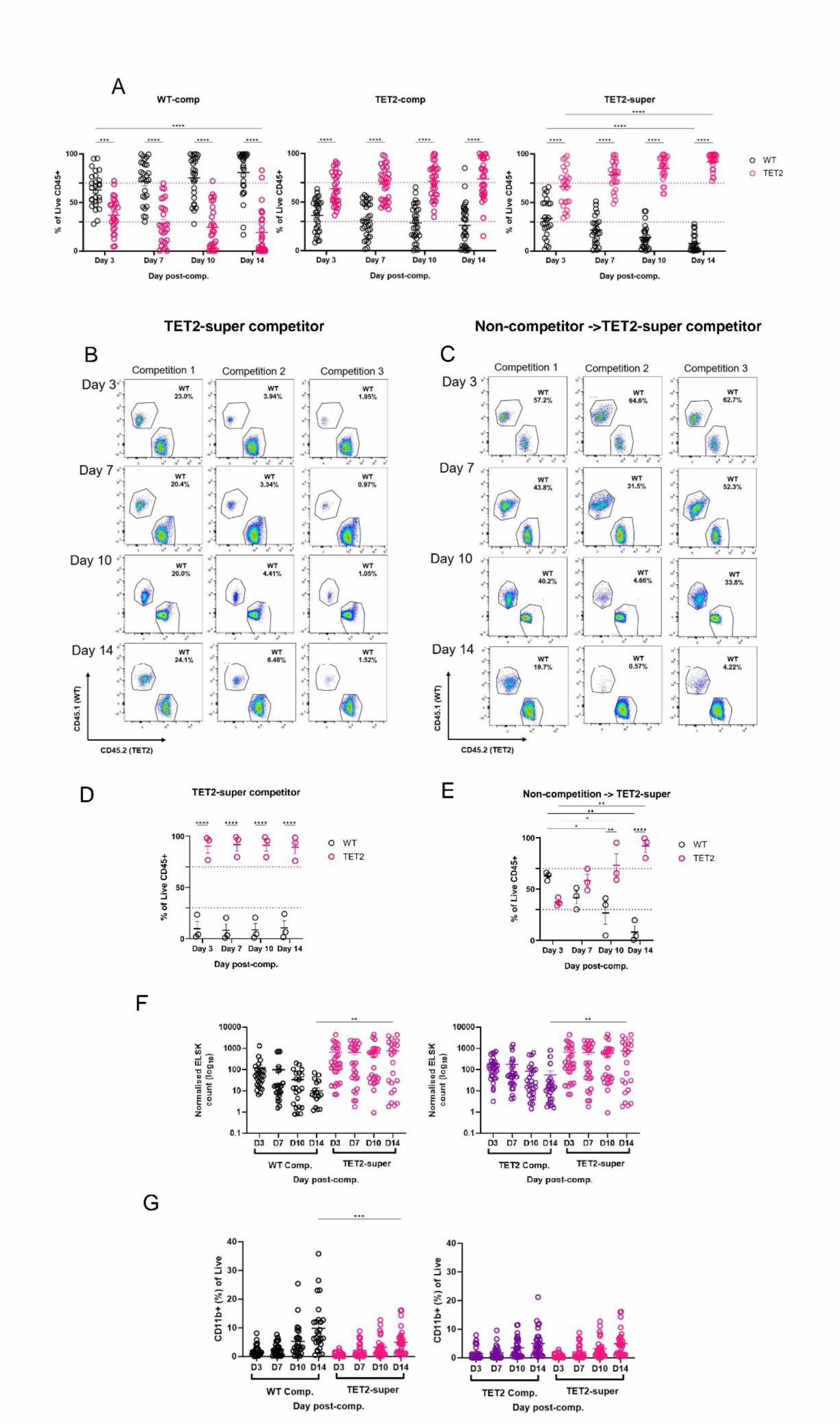
Temporal phenotypic analysis of wild-type and TET2 competition cultures on days 3, 7, 10 and 14 of the competition experiment. **A)** Frequencies of Live CD45^+^ cells in competition cultures where wild-type (left), TET2 (centre) and TET2-super (right) competitor clones were found, throughout the duration of the competition experiment. Dominance of WT and TET2 competitors was exerted from the early timepoints of the culture and was stable throughout the experiment. TET2-super competitors, on the other hand, displayed more nuanced dynamics. **B-C)** Individual examples of individual TET2-super competitor clones and their dynamics through time. **B)** Example of a clone which exhibited a stable, super-competitive phenotype through time. These clones represented 25% of TET2-super competitors in this experiment. **C)** Example of a TET2 super competitor clone which had no initial advantage, but acquired the super-competitor phenotype over time. These clones represented 25% of total TET2-super competitor clones in the experiment. Percentages of CD45^+^ cells illustrate the competition dynamics. **D-E)** Dot plots representing the temporal dynamics of cell competition for each of the individual examples shown in **B-C**. **F-G)** Comparisons of ELSK **(F),** and CD11b^+^ **(G)** frequencies between different competitor types throughout time. Accordingly, TET2-super competitors had the highest frequency of ELSK and LSK cells across time and across competitor types. In contrast, WT-competitors displayed the highest frequency of CD11b^+^ monocytes at the end of the competition experiment (Day 14) in comparison to TET2-super competitors. Two-way ANOVA, * p < 0.05, ** p < 0.01, *** p < 0.001, **** p < 0.0001. Error bars represent the standard error of the mean (SEM).

We next analyzed cell populations by flow cytometry in WT competitors, TET2 competitors, and TET2-super competitors (**Figure 3F-G** and **Supplementary Figure 3C-F**). Consistent with our previous findings, TET2-super competitors had elevated HSPC numbers by day 28 (**Figure 3F**), which was also evident across time (3 days post-split and cross-competition), as well as in the unmixed controls (**Supplementary Figure 3E**). ELSK and LSK absolute numbers, on the other hand, remained at comparable levels for the WT and TET2 competitors (**Supplementary Figure 3C**). In contrast, WT competitors contained a higher proportion of CD11b^+^ cells (**Figure 3G** and **Supplementary Figure 3G**). We also compared mature cells in the sister WT controls of suppressed clones (i.e., WT clones that “lost” to the TET2-super competitor clones) and found that higher numbers of CD11b^+^ cells in the control sister wells correlated with poor performance of the sister well against TET2 super-competitor cells (**Supplementary Figure 3H**).

### Non-intrinsic mechanisms mediate super competitive phenotype

We next interrogated competitor clone functional potential via *in vivo* transplantation. Control wells corresponding to the sister wells defined as WT competitors, TET2 competitors, or TET2-super competitors were harvested, and 30% of cells were transplanted into recipient animals at the experimental endpoint of the competition assay (14 days post-split). At 16-weeks post-transplantation, only TET2-super competitors had a 100% repopulation rate (**Figure 4A**) and both TET2 and TET2-super competitor cell recipients displayed higher donor-derived chimerism (**Figure 4B**) compared to WT competitors, mirroring the increased number of phenotypic HSCs observed in competition cultures (**Figure 3F**). Subgroups of good and bad engrafters were observed in TET2 competitor transplant recipients, however, these exhibited the highest chimaerism levels among competitor clone types. All competitor types were capable of multilineage reconstitution (**Supplementary Figure 4A-B**). While clones with higher phenotypic HSCs in TET2 competitor cultures had a strong positive correlation with functional output (r^2^ = 0.6048, p = 0.00396) (**Figure 4C**), this was not observed for TET2-super competitor or WT-competitor clones (**Supplementary Figure 4C**). This latter finding would be consistent with the most effective TET2-super competitor clones not giving rise to as many mature cells, in accordance with the long-term differentiation impairment reported in *Dnmt3a-*null HSCs (Challen et al, 2011; Jeong et al, 2018). We also analysed BM and spleen of recipients at 24 weeks post transplantation and found that TET2-competitor clones had the highest fraction of donor-derived CD45^+^ cells (**Figure 4E**), largely in agreement with chimerism levels in circulating white blood cells and LSK fractions in both tissues (**Supplementary Figure 4D**). As it remains possible that TET2-super competitors do not simply have more HSC functional potential, this prompted us to next investigate the non-HSC fraction of TET2-super competitors for their capacity to suppress WT cells.

**Figure 4.**
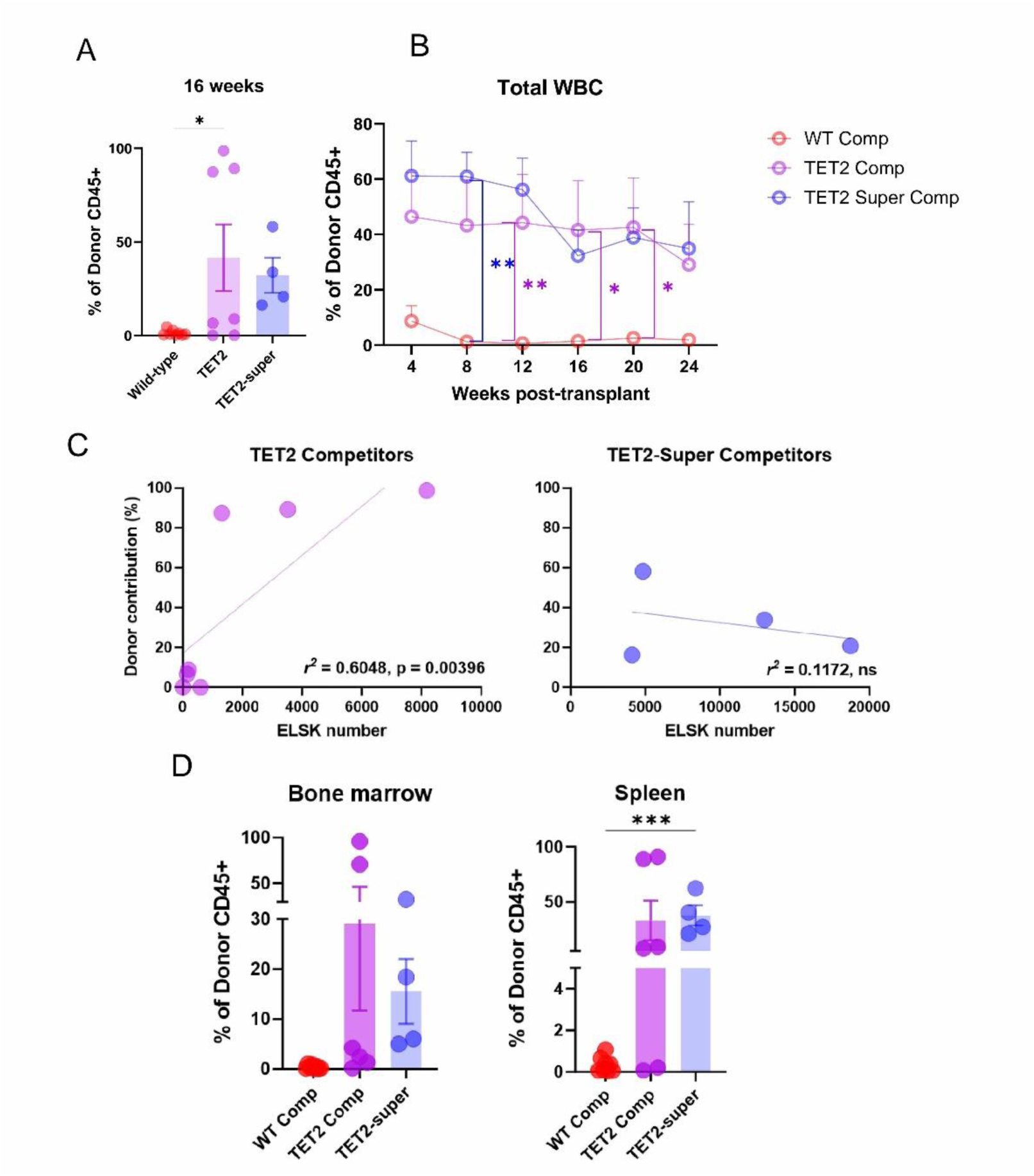
*In vivo* functional characterization of competitor types. **A-B)** Analysis of donor contribution to total WBC fraction in recipients of WT- (n = 8), TET2- (n = 8), and TET2-super-(n = 4) competitor clones, from 4-24 weeks post-transplant. **A)** Donor-derived chimerism at 16 weeks post-transplant. For *in vivo* transplantations, 30% of corresponding, non-competed controls was harvested, and injected into recipients together with bone marrow supportive cells (see Supplemental Methods). WT-competitors had repopulated 62.5% of recipients, TET2-competitors 71.42% of recipients. In contrast, all TET2-super competitor clone recipients had been repopulated by 16 weeks. Bars represent mean and error bars represent standard error of the mean (SEM). **B)** Donor-derived chimerism throughout the experiment. TET2 competitors attained the highest chimaerism levels, followed by TET2-super competitors, and lastly, by WT competitors. Two of the TET2-competitor recipients died before the completion of the experiment due to reasons unrelated to the transplant. **C)** Linear regression analysis of ELSK absolute numbers in relation to donor-derived chimerism in TET2- (left) and TET2-super competitors. ELSK counts *in vitro* significantly correlated with donor-derived chimerism *in vivo* for TET2-competitors (r^2^ = 0.6048, p = 0.00396). Average ELSK number in TET2-competitor donors (μ = 1.993 x 10^3^) was ten-fold reduced relative to TET2-super competitors (μ = 2.664 x 10^4^). **D)** Competitor clone overall contribution to the live CD45+ fraction in the bone marrow (left) and spleen (right) of recipients. A) One-way ANOVA; B), Two-way ANOVA; C) Simple linear regression model; D) One-way ANOVA, * p < 0.05, ** p < 0.01, *** p < 0.001, **** p < 0.0001. Error bars represent the standard error of the mean (SEM).

### Upregulation of inflammatory pathways and cell adhesion molecules in TET2-super competitors

To decipher the potential molecular drivers of the TET2-super competition, we profiled competitor clones by RNA-sequencing. Principal component analysis did not separate competitor phenotypes into distinct clusters (**Supplementary Figure 5A**), so we filtered the globally top 250 expressed genes across all competitor clones (**Supplementary Figure 5B and Supplementary Table 1**) to identify TET2-super competitor specific changes. These were fundamentally characterized by elevated expression of cytoskeletal genes and inflammatory pathways, including *Abca6, Lad1,* and *Tlr9*, which have previously been implicated as modifiers of cell outgrowth in solid tumours (Hedditch et al, 2014; Hu et al, 2023) and leukemia (Kennedy et al, 2021). Differential gene expression analysis by DESeq2 (Love et al, 2014) (**Figure 5A**, **Supplementary Table 2**) identified structurally relevant genes including *Adgrg5,* an adhesion G-protein coupled receptor (GPCR) (Wilde et al, 2016; Liessmann et al, 2024), *Dock4,* a guanine exchange factor (GEF), a member of the Rho-GTPase family (Hiramoto et al, 2006; Ueda et al, 2013), and CYFIP related Rac1 interactor A (*Cyria*) (Fort et al, 2018). KEGG enrichment analysis (**Figure 5B, Supplementary Table 3**) mainly included response to interferon (IFN)-β, regulation of viral infection, and negative regulation of viral processes. Specifically, the IFN-induced genes, *Ifi206* and *Ifi208* contributed to many enriched terms, pointing to a dysregulated inflammatory response in TET2-super competitor clones and further according with elevated interferon gene expression and dysregulated IFN-γ described in mouse models (Xie et al, 2023; Jenkins 2025). Collectively, altered genes pointed to dysregulation of both RHO-GTPases and inflammatory pathways upon TET2-super-competition (**Figure 5C**).

**Figure 5.**
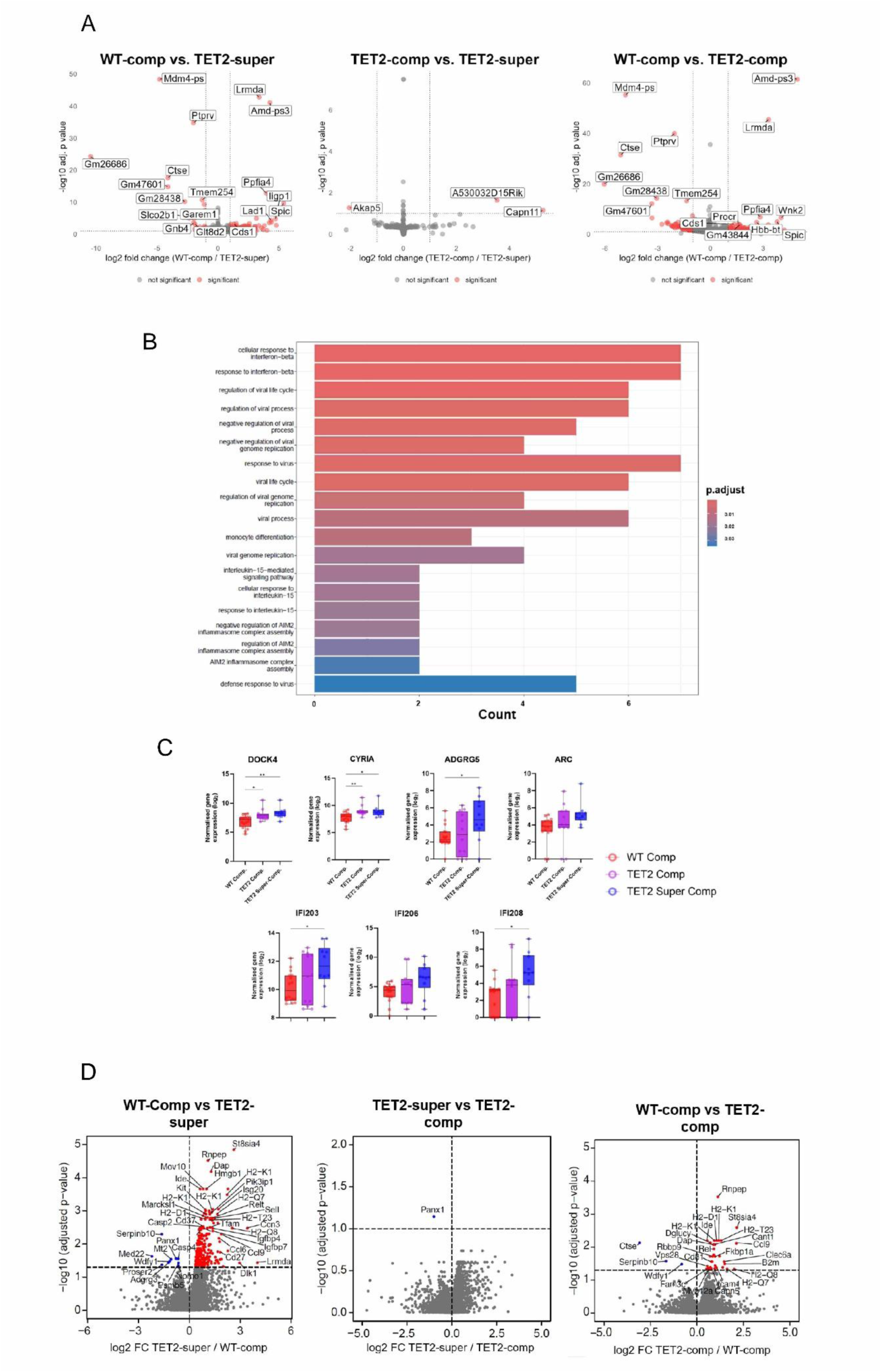
TET2-super competitors have elevated inflammatory and cell adhesion-related molecules. **A)** Molecular profiling of competitor clones. A total of 30 clones (17 WT, 13 TET2, and 10 TET2-super competitor clones) were analysed, obtained from three independent experiments. Differential gene expression (DGE) analysis was performed in pairwise comparisons: WT-vs TET2-super-, WT- vs TET2-, TET2-super- vs TET2-competitors. A total of 38 clones were analysed across competitor types (WT-comp n = 16, TET2-comp n = 12, TET2-super n = 10). Volcano plots represent each of these comparisons. **B)** Gene ontology (GO) enrichment analysis was performed in pairwise comparisons between WT- and TET2-super, and TET2- vs TET2-super-competitor clones. In the first comparison, 19 pathways were found to be significantly enriched. Cellular response to IFN-β (p = 8.64e-08) and response to IFN-β (p = 1.33e-07) were the main upregulated pathways in TET2-super competitors. **C)** Individual gene expression analysis for molecules implicated in cytoskeletal organisation, *Dock4* and *Cyria*, cell adhesion *Adgrg5* and *Arc* and interferon-induced proteins *Ifi203, 206* and *208*. In addition, we also observed genes implicated in immune cell regulation and function, including *Cd160, Cd4, Fcer2a,* and *Cd209a*, H2-Q6, and H2-Q7 as well as genes associated with ion binding activity like *Efcab7* and *Scel* (Cui et al, 2024; Champliaud et al, 1998).Gene expression data was normalised using the median of ratios method in DESeq2 **D)** Differential protein abundance analysis of WT- (n = 14), TET2- (n = 18), and TET2-super- (n = 16) competitors (total n = 48), from two late culture timepoints (days 26 and 28), totalling 51 clones analysed. Pairwise comparisons were performed in a similar manner as for DGE. Analysis of the secretome by DeepLoc protein localisation (Ødum et al, 2024) validated the molecular DGE and GO enrichment analyses, wherein inflammatory molecules were found to be upregulated in TET2-super competitors. Count matrices were normalised and log-transformed (log_10_). One-way ANOVA, * p < 0.05, ** p < 0.01. Error bars represent standard deviation (SD).

We also analysed the global secretome of different competitor clone subtypes (**Figure 5D**). Here, we identified 234 proteins with differential abundance in the TET2-super competitors compared to WT competitors (adjusted p<0.05, log_2_FC >0.5) (**Figure 5D**), 75 of which were classified as probable secreted proteins by the DeepLoc protein localization tool (Ødum et al, 2024) (**Supplementary Table 4**). Interferon signalling was again increased, including IFI35 and IFNAR2, and a series of different MHC-II complex members. Together, our transcriptomic and secretomic profiling data converge on the upregulation of inflammatory pathways and cell adhesion-related molecules typifying super-competitive TET2 clones but whether they might play a functional role in suppressing WT cells remained unclear.

### Preleukemic cell resilience to RAC1/GEF inhibition

To find specific molecules that selectively target non-mutant cells while sparing mutant cells, we undertook targeted functional assays on a range of targets, including interferon and immune cell signalling (**Figure 6A**). Viability of WT cultures, and to a lesser extent TET2^-/-^ cells was significantly reduced by addition IFN-γ in a dose-dependent manner (**Supplementary Figure 6A**), demonstrating that while higher concentrations of IFN-γ are indeed toxic to WT cells, they are also toxic to TET2^-/-^ cells and therefore not selective enough to drive the super-competitor phenotype.

**Figure 6.**
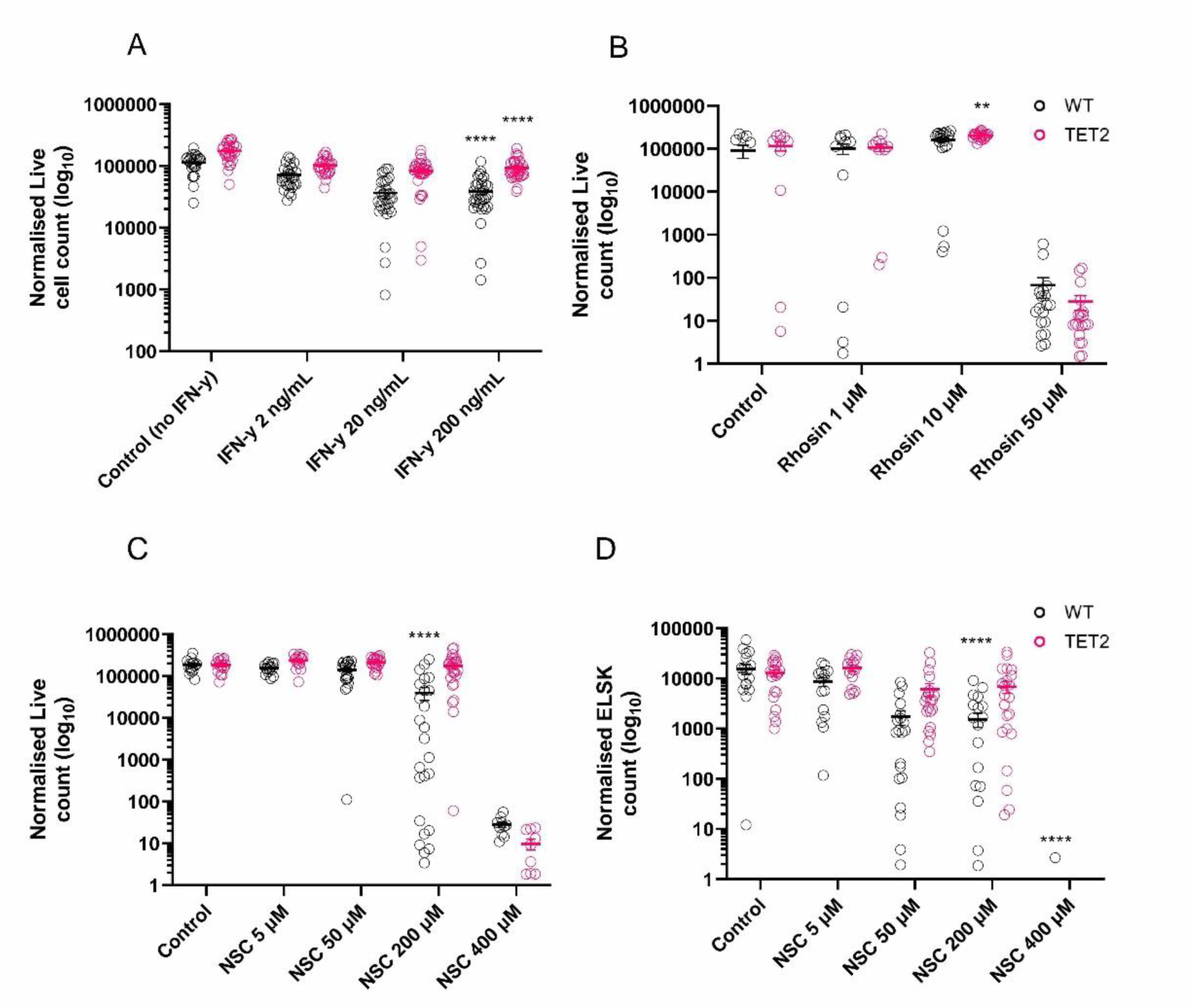
Disruption of Rac1/GEF interaction is selectively tolerated by TET2^-/-^ HSCs. Bulk HSCs were cultured and treated with different molecules at different doses. Comparisons were made against the matched untreated control for each genotype. **A)** Treatment with IFN-γ caused a significant reduction both WT and TET2 cells, especially at the highest dose. **B)** Rho-GTPase signalling was disrupted by treatment with Rhosin and led to toxicity in both WT and TET2^-/-^ cells. **C)** NSC-23766, a specific inhibitor of Rac1/GEF interaction, selectively reduced WT cell viability while maintaining TET2 viability up to 200 μM. **D)** ELSK numbers in wild-type cultures were likewise reduced in a dose-dependent manner, in relation to the control (non-treated) conditions. TET2 ELSK number, in contrast, was not significantly affected by NSC-23766 treatment up to 200 μM concentration. Two-way ANOVA, * p < 0.05, *** p < 0.001, **** p< 0.0001. Error bars represent the SEM.

We next investigated the effect of Rho-GTPase signalling disruption during WT and TET2-mutant HSC expansion. Given our finding of elevated expression of *CYRIA*, which encodes for the Rac1 member of the Rho-GTPase family, along with the GEF, DOCK4, in TET2-super competitor clones, we hypothesised that upregulated Rho-GTPase signalling supported super-competitive behaviour. We tested three different molecules, which were either selective for specific Rho-GTPase members, or pan Rho-GTPase inhibitors. Treatment of WT and TET2 cultures with CASIN (a selective CDC42 inhibitor) and Rhosin (pan Rho-GTPase inhibitor) were toxic (**Supplementary Figure 6B-C**). Treatment with an inhibitor to the third major subfamily of Rho-GTPases, Rac1 GTPases (Cancelas et al, 2009; Cancelas et al, 2011) via NSC-23766, a Rac1/GEF interaction inhibitor considered to be CDC42-independent (Gao et al, 2004), however, showed a striking and selective effect. While very high doses (400 μM) were toxic to both WT and TET2^-/-^ cells, both the 200μM and 50μM doses showed a strong selective, negative impact on WT cells (**Figure 6C**). The 200 μM dose resulted in 4.42-fold fewer WT cells compared with TET2^-/-^ cultures and the total number of phenotypic HSC was likewise significantly reduced by 4.45-fold (**Figure 6D**, **Supplementary Figure 6D)**. Together, these data identify inhibition of the Rac1/GEF interaction as a selective mechanism for TET2 mutant cells to establish a toxic environment for WT cells as an early step for gaining a competitive advantage in their local microenvironment. These data also demonstrate more generally that this clonal competition assay is a powerful *in vitro* model that helps identify novel single-clone mediated effects that can modulate cell competition dynamics.

## Discussion

In this work, we have developed and optimized a split culture method that can be leveraged for *in vitro* modelling of cell competition at clonal resolution. Our approach offers: 1) clonal resolution; 2) scalability; and 3) direct linkage to functional outputs. Similar approaches, including paired daughter cell assays (Suda et al, 1984; Suda et al, 1984; Leary et al, 1985; Ema et al, 2005; Kent et al, 2008; Weinreb et al, 2020; Wehling et al, 2022) and HSPC split cultures (Benveniste et al, 2010; Naik et al, 2013; Lin et al, 2018; Tian et al, 2021) have been restricted to clonal tracking for describing cellular or molecular outputs, assuming substantial uniformity of cell output in daughter HSPCs. Our approach, which splits HSC clones into phenotypically, functionally and molecularly equivalent fractions, provides direct control comparators for simultaneously delivered perturbations. We increase scalability with 1,260 competition wells analysed across all experiments, corresponding to >300 individually profiled clones. A particular advantage is the ability to cross-compete clones of different genotypes to explore their relative competitive ability, leading to the direct discovery of a novel preleukemic mechanism of WT cell suppression.

In this study, we identify and describe TET2-super competitor clones which are present in all experiments conducted and actively suppress the growth of WT cells. Super-competition has been described in stem cell systems whereby the stem cell population enhances its proliferation at the expense of other clones (Di Gregorio et al, 2016; Yusupova & Fuchs, 2023; Cong & Cagan, 2024; Perez Montero et al, 2024). Moreover, previous reports have provided global evidence for the competitive advantage of TET2-mutant clones (Yang et al, 2025), as well as DNMT3A, ASXL1 (Cooper & Young, 2017) and other hematological mutations in relation to non-mutant clones (Ferrando & Lopez-Otin, 2017; Watson et al, 2024), but studying these processes at the single-cell level has been severely limited. In addition, cell heterogeneity across the hematopoietic differentiation spectrum hinders the ability to discern molecular differences in an *ex vivo* setting (Singh et al, 2025). Split culture followed by clonal competition provides a scalable platform for studying this phenomenon in HSCs.

Our molecular and secretomic studies inferred a TET2-mediated upregulation of interferon response pathways and cell adhesion molecules. The interferon pathway, particularly IFN-γ, accords with previous data (Marino et al, 2016; Cheng et al, 2024; Herbrich et al, 2025; Heavican-Foral et al, 2026), but our functional studies did not reveal genotype-specific differences in expansion cultures. This nevertheless warrants further testing to find a nuanced mechanism by which inflammatory pathways could potentially enable super-competitive behaviour. On the other hand, the identification of cell adhesion molecules in super-competitor TET2 cells complements our previous work which identified distinct extracellular matrix regulation in TET2-mutant HSCs (Jassinskaja et al, 2026) and led us to pursue Rho GTPase signaling cascades as potential mediators. While CDC42 and RHOA inhibition negatively impacted both WT and TET2-mutant cells, the CDC42-independent Rac1 Rho-GTPase family inhibitor NSC-23766 was shown to be a selective mediator of WT cell suppression. Both CDC42 (Florian et al, 2012) and RHOA (Xu et al, 2009), have been previously implicated in HSC function, perhaps explaining their uniform negative impact on HSC expansion in culture. Rac1 GTPases therefore represent a WT-specific vulnerability with previously implicated roles in physiological cell migration (Fort et al, 2018; Le et al, 2021) and disease-related inflammation (Zang et al, 2017). Interestingly, off-target effects of NSC-23766 have been identified on platelets in a Rac1^-/-^ setting (Dutting et al, 2015), which might impact modulatory roles of platelets and megakaryocytes during HSC expansion. Considering its highly selective effect between WT and TET2 mutant cells, Rac1 Rho-GTPase signalling could therefore be harnessed both for the promotion of HSC expansion or for slowing/preventing the early stages of leukemic clone establishment *in vivo*.

Overall, this work introduces split clone cell competition assays as a powerful tool for studying the earliest stages of establishing clonal dominance by single CH-mutation bearing HSCs. Cultures can be split into homogeneous fractions, and these can be cross-competed (or cross-treated) to enable discovery of novel subsets of cells with clinical significance or novel therapeutics that specifically modulate the clonal expansion dynamics of cells with specific mutations. Here, we have used this system to describe an interesting cell phenotype and corresponding behaviour - a single cell-initiated clone that can suppress multiple WT counterparts simultaneously, creating a positive feedback loop that enables clonal dominance to be established, with CDC42-independent RAC1 signaling as a central and selective mediator. This *in vitro* platform is thereby widely applicable, and offers increased statistical power to conduct clonal competition experiments at scale. Not only could these competitor cells and their respective molecules be considered as novel therapeutic targets, but these studies also point toward a greater need to study both the initiating cell and its progeny in concert, giving us a template for modelling more complex disease paradigms.

## Data availability

sc-RNA sequencing data for single-cell split cultures and bulk RNA-sequencing data for competitor clones have been deposited in the ArrayExpress controlled access repository (E-MTAB-17115 and E-MTAB-17274, respectively).

Mass spectrometry data and proteomics results files for competitor clones have been deposited in MassIVE (MSV000101882) and referenced in ProteomeXchange (PXD078551).

Sequencing and proteomics data were deposited under the study title: “A scalable in vitro cell competition model identifies preleukemic super competitor clones”.

## Acknowledgments

The authors would like to thank Graeme Park, Sukhveer Mann, Karen Hodgkinson, Samantha Donninger, Lesley Gilbert, Chris Taylor, and Peter O’Toole from the Technology Facility at the University of York for their support throughout this work. We likewise thank members of the York Centre of Excellence in Mass Spectrometry, which was created thanks to a major capital investment through Science City York, supported by Yorkshire Forward with fund from the Northern Way Initiative, and subsequent support from EPSRC (EP/K039660/1; EP/M028127/1). We warmly thank Rebeccah Hannah and Karsten Bach from the University of Cambridge for assistance with the Parse Bioscience single-cell pipeline. Thank you to Katarzyna Kania from the Genomics Unit at the University of Cambridge for processing and troubleshooting of Parse libraries. We especially acknowledge all members of the Kent Laboratory and the Centre for Blood Research, particularly Ian Hitchcock and Dawn Coverley for their support and helpful discussions. We would also like to extend our thanks to the team at the Biological Services Facility of the University of York, who have been an essential part of this work. A very special thank you to Dave Boucher and Simona Valletta for valuable advice and constructive feedback.

Work in the Kent laboratory has been supported by European Research Council Starting Grant (ERC-2016-STG-715371), a Cancer Research UK Programme Foundation Award (DCRPGF\100008), an MRC-AMED joint award (MR/V005502/1), the UK Medical Research Council (MC_PC_21043; MR/Y011945/1), Blood Cancer UK (24014), and the Swedish Research Council (to M.J.: 2021-00185). The Kent Laboratory is also supported by the National Institute for Care and Health Research Leeds Biomedical Research Centre (NIHR203331). Work in the Göttgens laboratory is supported by Wellcome (206328/Z/17/Z and 309075/Z/24/Z), Blood Cancer UK (18002), Cancer Research UK (C1163/A21762), MRC-AMED joint award (MR/V005502/1), MRC (MR/W031663/1), Blood Cancer United (7035-24) and Aging Biology Foundation. This research was funded in whole, or in part, by the Wellcome Trust [203151/Z/16/Z, 203151/A/16/Z and 226795/Z/22/Z] and the UKRI Medical Research Council [MC_PC_17230]. For the purpose of open access, the author has applied a CC BY public copyright licence to any Author Accepted Manuscript version arising from this submission.

## Authorship

L.C.C., J.A.R.L., N.K.W., and D.G.K. conceived of the project and designed the experiments. L.C.C., J.A.R.L., E.B., and J.M. performed competition experiments, harvested samples, performed flow cytometry, RNA extraction and *in vivo* transplantations. L.C.C. J.A.R.L. and K.H. performed single-cell and bulk-cell isolation. S.J. and S.M. performed RNA-sequencing experiments and data processing, respectively. L.C.C. performed bulk RNA-seq analysis with S.M., under the supervision of D.G.K. J.A.R.L. and M.S.V performed scRNA-seq analysis under the supervision of N.K.W. and B.G. M.J., C.B. and L.C.C. designed secretomic profiling experiments under the supervision of A.D. and D.G.K. L.C.C., C.B., and M.J. performed DIA-MS experiments, and M.J. analysed DIA-MS data. L.C.C. and D.G.K. wrote the manuscript with input from all authors. All authors read and approved the manuscript.

## Supplementary Figures

**Supplementary Figure 1.**
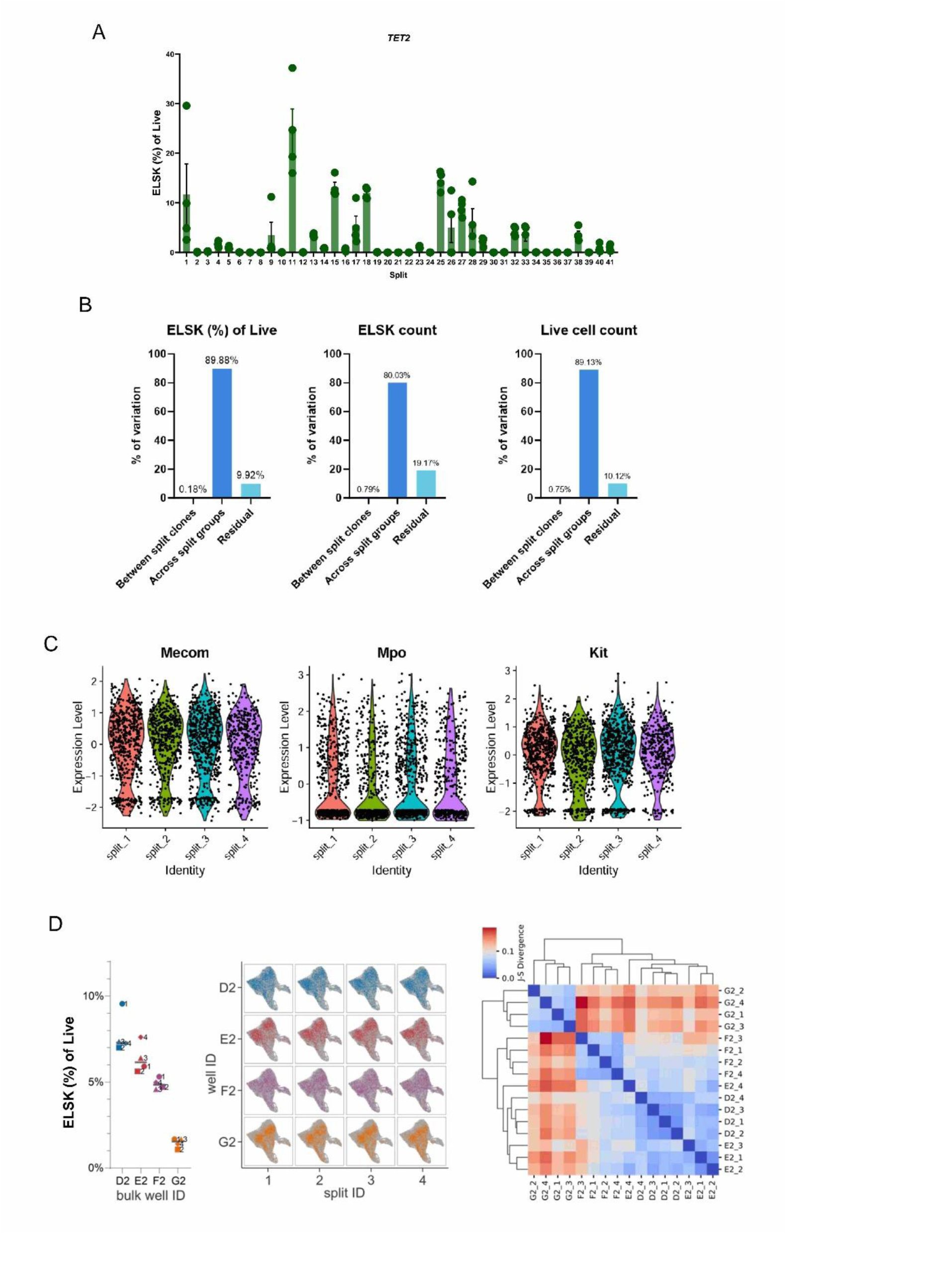
Extended phenotypic and molecular analysis of split split cultures. **A)** ELSK frequencies calculated from live cells for TET2 split cultures (n = 41). TET2 single-cell derived clones were split by the same method described for wild-type clones. Bars represent mean values; error bars represent the SEM. Variance partitioning analysis to understand variation in ELSK frequency of live, ELSK counts, and viable cell counts across and split clonal fractions. In each instance, differences in frequencies or counts came from across split groups (n = 38), rather than from within split clonal fractions. In agreement with ICC analysis (Koo & Li, 2016), this was evidence for the phenotypic consistency within split cultures. Bars represent percentage of data variation according to ICC model 3 (two-way mixed model). **C)** Individual gene expression of *Mecom*,*Mpo* and *Kit* across subclones. In a similar way to global gene expression, expression of genes implicated in HSC self-renewal was homogeneous across split cultures. **D)** ELSK frequencies (left), UMAP profiles (centre), and heatmap (right), representing the phenotypic and transcriptomic equivalency of four replicate bulk split cultures. Bars represent the mean and error bars represent standard deviation.

**Supplementary Figure 2.**
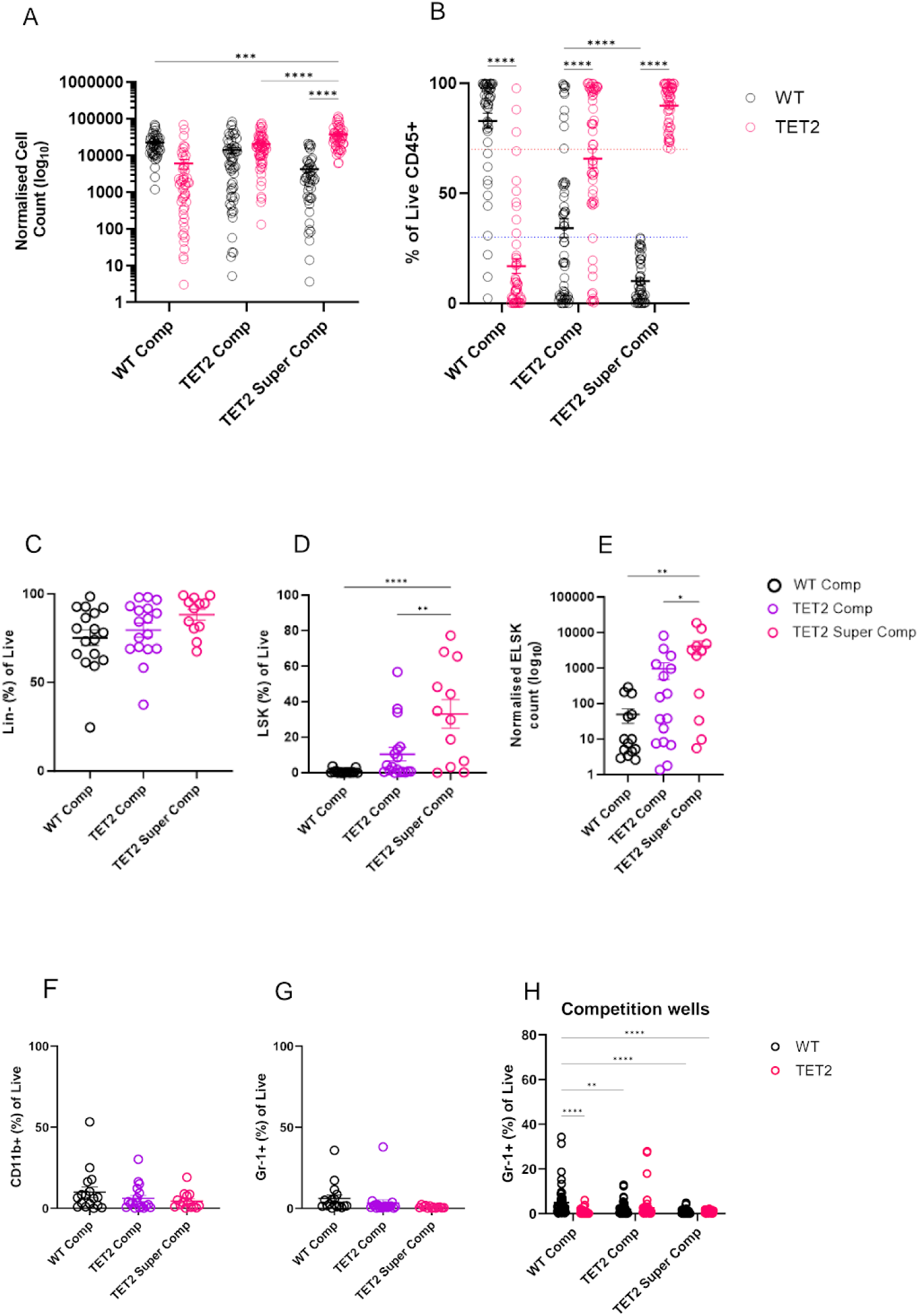
Extended phenotypic profiling of competitor clones. **A)** Normalised cell count and **B)** frequencies of live CD45+ cells in competition wells for each competitor types. **C-H)** Phenotypic analysis in control (non-competed) clones corresponding to competitor clones in **(A)** and **(B)**. **C)** Lin- and **D)** LSK frequencies, and **E)** ELSK numbers. Lin-frequencies were comparable across competitors. However, LSK and ELSK fractions were significantly elevated in TET2-super competitor clones. **F)** Frequencies of CD11b^+^ monocytes and **G)** Gr-1^+^ granulocytes were not significantly elevated for any of the competitor types. H) In competition wells, Gr-1+ cells were significantly elevated in WT-competitors. A-B, H) Two-way ANOVA, C-G) One-way ANOVA, * p < 0.05, ** p < 0.01, *** p < 0.001, **** p< 0.0001. Error bars represent the SEM.

**Supplementary Figure 3.**
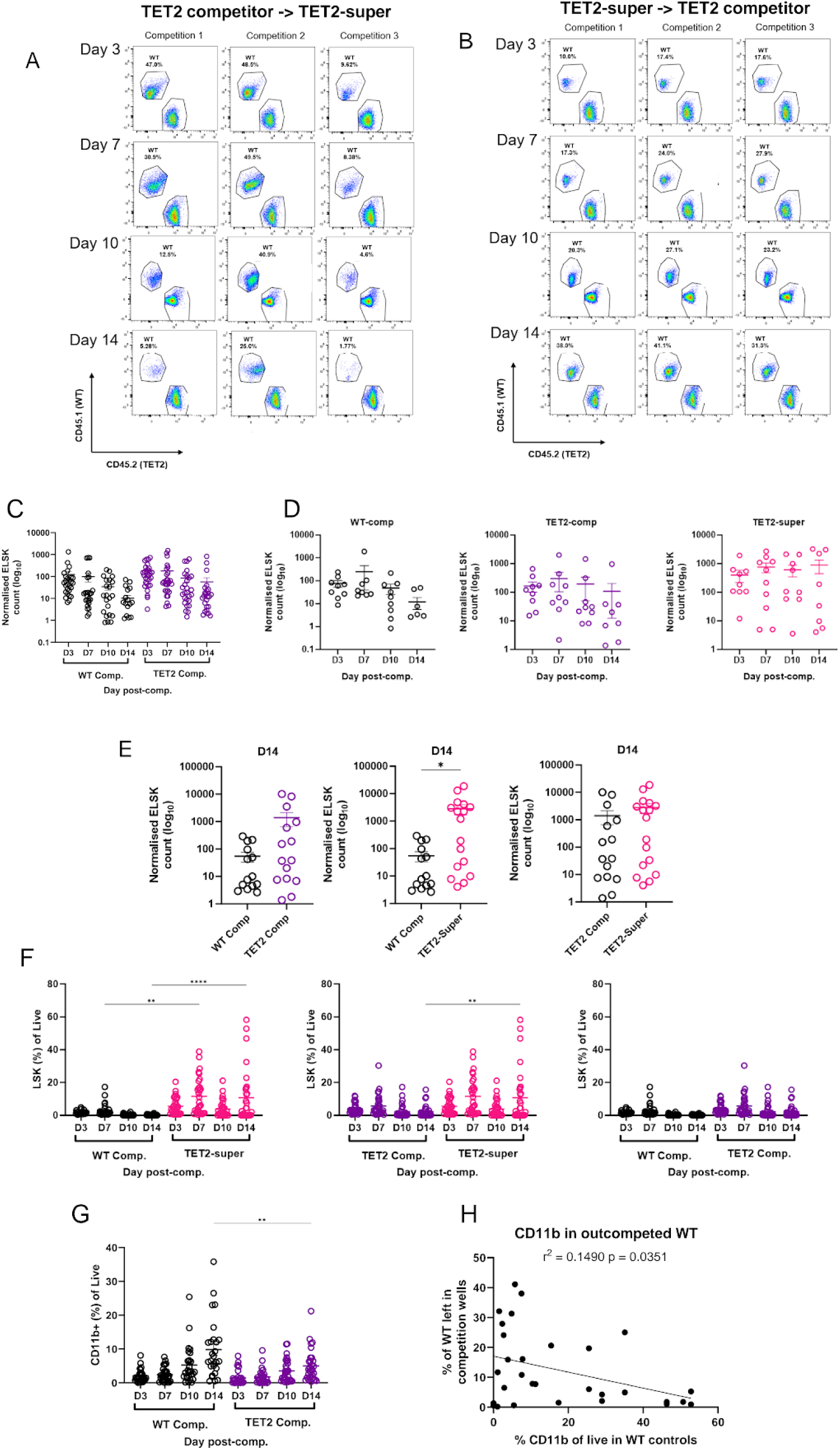
Extended temporal dynamics profiling of competitor clones. **A)** A representative example of phenotype transition from TET2- to TET2-super competitor through time. These clones represented 50% of total TET2-super competitors in this experiment. **B)** Representative example of a clone which displayed a seemingly TET2-super competitor phenotype, but was defined as a TET2-competitor due to its lack of dominance over wild-type rivals across a matrix. These clones were not considered as TET2-super competitors for the subsequent analyses. **C)** Comparison of ELSK numbers in WT- and TET2-competitors across time. **D)** Normalised ELSK counts in WT- (left), TET2- (middle) and TET2-super- (right) non-competed clones across time. Notably, ELSK content in TET2 non-competed controls was gradually, stably elevated from day 3 to day 14. **E)** Pairwise comparisons of ELSK numbers in control clones corresponding to each competitor type on day 14 of the competition experiment (28-day endpoint). In accordance with our findings, TET2-super control clones displayed a significantly elevated number of ELSK cells in comparison to WT-competitor controls. **F)** LSK proportions calculated from total live cells in pairwise comparisons across different competitor types. Accordingly, TET2- super competitors were shown to carry the highest percentages of LSK cells on day 14 of the competition experiment. **G)** CD11b+ percentages calculated from total live cells in WT-competitors were significantly elevated in comparison to TET2-competitors on day 14 of the experiment. **H)** Linear regression analysis showed that CD11b+ monocytes are elevated in WT non-competed controls corresponding to clones which were defeated by TET2-super competitor clones (r^2^ = 0.1490, p = 0.0351). One-way ANOVA, * p < 0.05, ** p < 0.01, *** p < 0.001, **** p< 0.0001. Error bars represent the SEM.

**Supplementary Figure 4.**
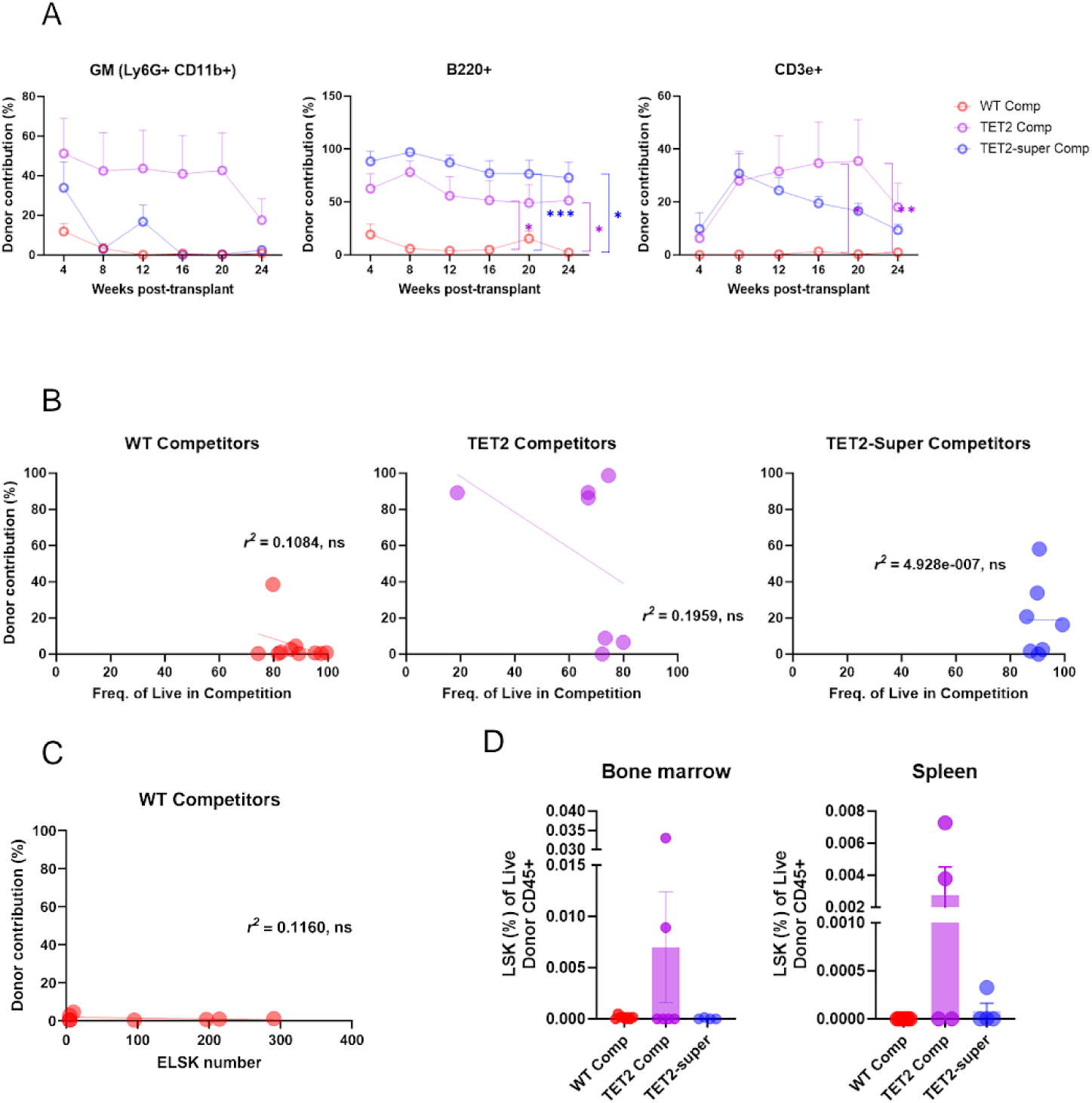
Extended functional profiling of competitor clones. **A)** Multilineage reconstitution analysis of competitor clones.Granulocyte-monocyte (GM) fraction percentages were higher in TET2-competitor recipients, albeit statistically non-significant, which agrees with their overall higher contribution to the total WBC fraction. Significantly elevated contributions to the B- (middle) and T-cell (right) lineages by both TET2- and TET2-super competitors provided evidence for their multilineage reconstitution capacity. **B)** Linear regression analysis of frequency of live CD45+ cells in competition wells relative to their donor-derived chimaerism levels. None of the correlations were statistically significant. **C)** Linear regression analysis of normalised ELSK counts with donor-derived chimaerism in WT-competitor recipients. **D)** Contribution to the LSK fraction in the bone marrow (left) and spleen (right) of recipients. TET2-competitors made higher contributions to the HSC/HSPC fraction in haematopoietic tissues, agreeing with their level of contribution to the overall circulating WBC fraction.

**Supplementary Figure 5.**
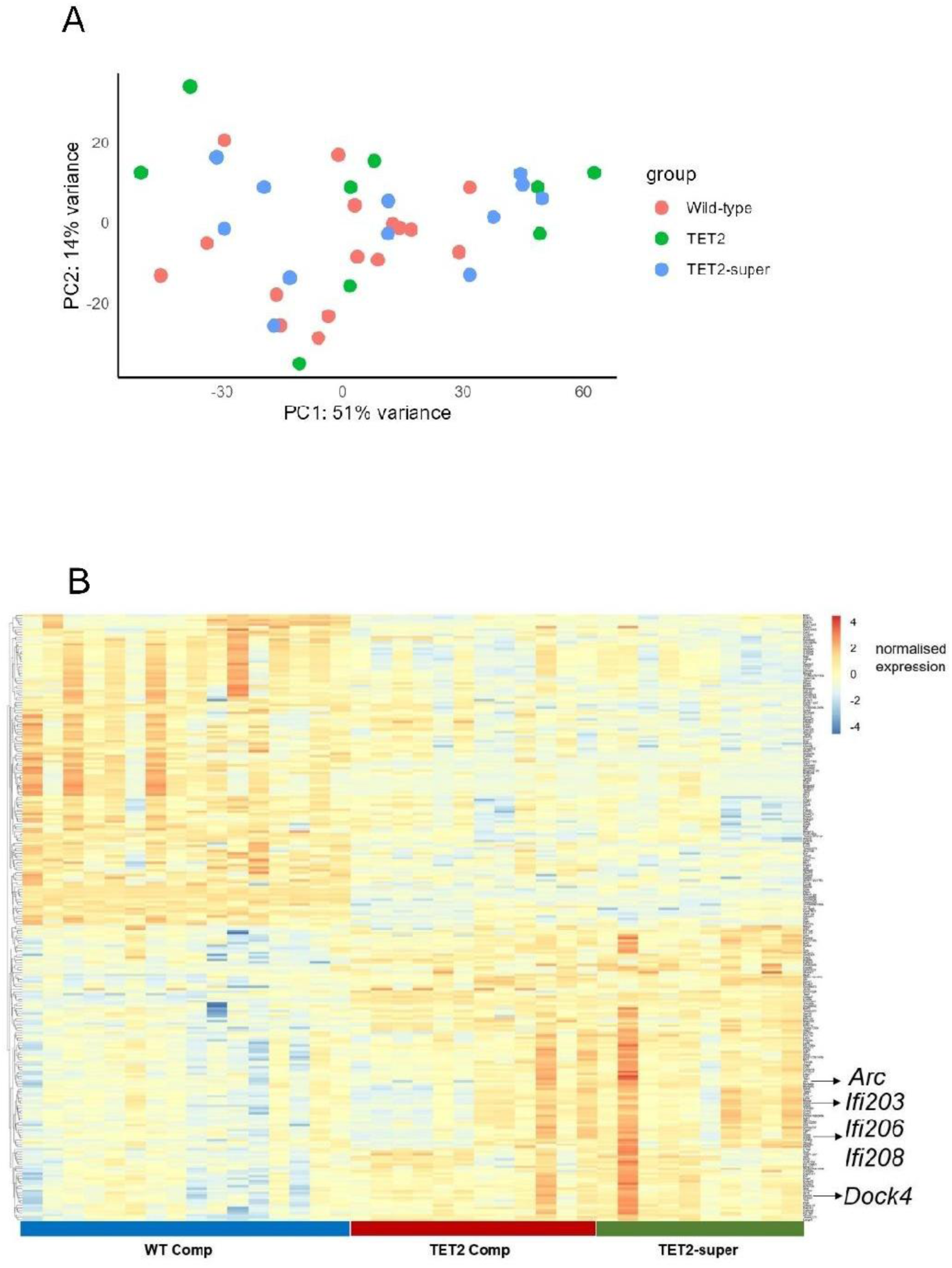
Extended molecular profiling of competitor clones. **A)** Principal component analysis (PCA) of wild-type-, TET2-, and TET2-super competitor clones. **B)** Heatmap of the top 200 expressed genes across competitor types. A signature of inflammatory, cell adhesion and cytoskeletal organisation molecules was detected in TET2-super competitors. Red-colour scale bar to the top right corner of the heatmap indicates relative gene expression.

**Supplementary Figure 6.**
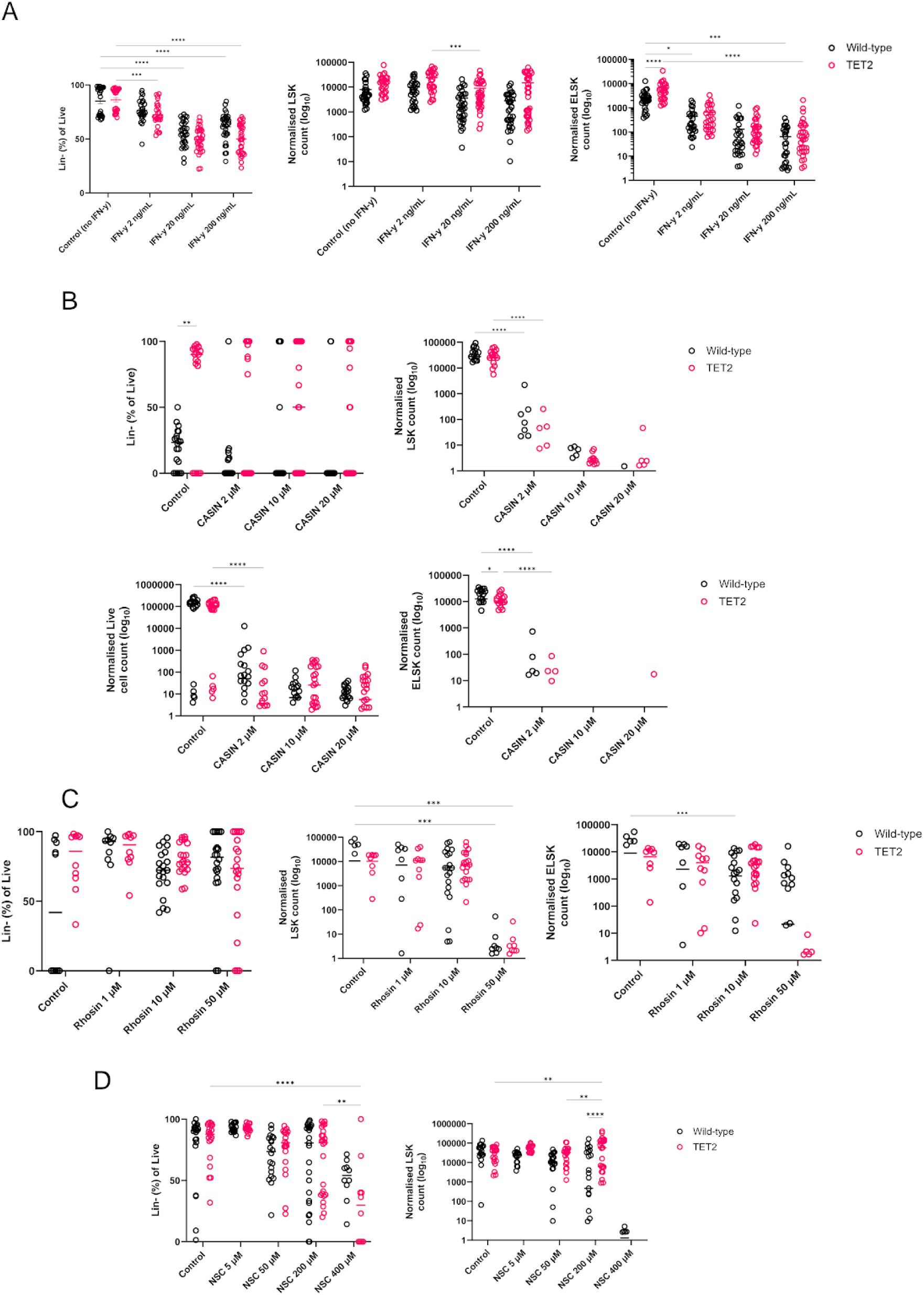
Extended phenotypic profiling of wild-type and TET2 bulk HSC cultures under different treatment conditions. Lin- frequencies calculated from live cells and LSK numbers are shown. **A)** IFN-γ treatment had a mild impact over the HSC/HSPC pool in both wild-type and TET2 cultures. **B)** As CASIN was lethal, cellular proportions were obliterated. No ELSK expansion was observed under these conditions. **C)** Rhosin negatively regulated the expansion of HSC/HSPCs in equal proportion for both genotypes. **D)** NSC-23766 treatment was tolerated by the HSC/HSPC fraction in TET2 cultures. A high dose (400 μM) diminished the proportions of these cells in both genotypes. Two-way ANOVA, ** p < 0.01, *** p < 0.001, **** p< 0.0001. Error bars represent the SEM.

