## Supplemental Methods for "A scalable *in vitro* cell competition model identifies preleukemic super competitor clones"

***Mice***

WT B6.SJL-*Ptprc^a^ Pep^c^/*BoyJ (hereafter, CD45.1) C57BL/6J (hereafter, CD45.2), *Tet2* loss-of-function B6(Cg)-*Tet2^tm1.2Rao^*/J (hereafter, TET2^-/-^) (Ko et al, 2011), and hybrid CD45.1/CD45.2 allele-harbouring (hereafter, hybrid) were bred and maintained in specific-pathogen-free facilities at the University of York. WT CD45.1 and CD45.2 mice were purchased from the Biological Services Facility (University of York, UK), TET2^-/-^ mice were obtained from Anjana Rao’s laboratory (La Jolla Institute for Immunology, US). Mice were continuously provided with sterile solid food pellets, water *ad libitum*, and bedding in microisolator cages. All regulated animal procedures were performed under Project License no. PEAD116C1 and PP9339632, approved by the Animal Welfare and Ethical Review Body (AWERB) of the University of York, and in compliance with the guidelines of operation of the Animal Scientific Procedures Act, 1986.

***Isolation of haematopoietic stem cells***

For single-cell or bulk-culture assays, femurs, tibias, hips, sternum and spine were collected from 8-12-week old WT (CD45.1) and TET2^-/-^ (CD45.2) mice and placed in PBS + 2% foetal bovine serum (FBS). Bones were crushed using pestle and mortar, and filtered through a 70-μm canister cell strainer to create a single-cell bone marrow suspension. Depletion of red blood cells was performed via incubation in ammonium chloride solution (StemCell Technologies) as previously described (Kent et al, 2009). For selective removal of the mature lineage fraction, bone marrow suspensions were incubated with a cocktail of biotinylated antibodies (StemCell Technologies) against CD3, CD5, CD11b, CD19, CD45R/B220, TER119, 7-4, and Ly6G/C, followed by immunomagnetic depletion with streptavidin-coated magnetic particles (StemCell Technologies). Enriched progenitor/stem cell fractions were stained with the antibody cocktail described in Supplementary Table 1, and incubated at 4^O^C for 30 min protected from light. Cell suspensions were filtered through a 40-μm cell strainer and resuspended into a 1:1000 7-Aminoactinomycin-D (Invitrogen) solution prepared in FACS buffer for the discrimination of viable from non-viable cells. HSCs were defined as EPCR^+^ CD45^+^ Sca-1^high^ CD150^+^ CD48^-^ (ESLAM-Sca-1^high^, hereafter, ESLAM) (Wilson et al, 2015; Che et al, 2022) were gated from total viable singlets (7-AAD^-^). Single or bulk (1 cell/ or 50 cells/well of a 96-well plate, respectively) ESLAM cells were isolated using the Moflo Astrios (Beckman Coulter) (equipped with 488-nm, 633-nm, 405-nm and 561-nm lasers) or the BD S8 FACS Discover (Waters Biosciences) 395-nm, 488-nm, 633-nm, 405-nm and 561-nm lasers) sorters. For single-cell assays, ESLAM cells were sorted in single-cell mode to ensure the deposition of one cell into each well; for bulk cultures, ESLAMs were sorted in purity mode.

***Ex vivo expansion of haematopoietic stem cells***

Clonal (single-cell) or bulk cultures in the F12-PVA expansion system were carried out as previously described (Wilkinson et al, 2020). Briefly, single HSCs were expanded in 96-well flat-bottom plates coated with 1mg/mL fibronectin from human plasma (Sigma Aldrich), each well containing 200 μL of Ham’s F12 nutrient mixture (Gibco), supplemented with 1% Penicillin/Streptomycin/L-glutamine solution (ThermoFisher), 1M Hepes buffer (Gibco), 1% Insulin-Transferrin-Selenium/Ethanolamine solution (Gibco), 10 mg/mL polyvinyl alcohol (PVA, Sigma Aldrich), 100 ng/mL animal-free, recombinant murine thrombopoietin (TPO, Peprotech), and 10 ng/mL animal-free, recombinant stem cell factor (SCF, Peprotech). Media was replaced with freshly prepared culture medium on a tri-weekly basis starting at day 5 or 7 after cell sorting. Cells were cultured inside a humidified incubator at 37^o^C, with 5% CO_2_ and 20% O_2_ levels for 14-28 days in total.

***Splitting of single-cell derived clones***

On day 14, single-HSC derived clones were visually assessed and viable cell numbers were calculated using Precision Counting Beads (Biolegend) and flow cytometry analysis. Cloning efficiency was estimated by dividing the number of clone-forming HSCs by the total numbers of wells sorted per plate. Total viable cells were estimated to be between 4.2x10^4^ - 3.3x10^5^ cells per clone. For splitting, contents of each well were mixed by gentle but thorough pipetting and each clone was distributed across four wells of a new plate (i.e., 25% of the original clone). Split sub-clones were carried for an additional 14 days (28 days total) and were subjected to downstream applications.

***Cell competition assays***

The CD45.1/CD45.2 mouse congenic system was used to distinguish murinw WT (CD45.1) from TET2^-/-^ (CD45.2) HSC-derived clones. WT and TET2^-/-^ HSCs were isolated and cultured separately for 14 days and successfully expanded clones were split and mixed in equal proportions in a *“Competition Matrix”*. WT clones were distributed horizontally and TET2^-/-^ cells vertically across the competition matrix, resulting in one control well per clone and three competition cultures per clone. Within the matrix, nine competition wells each contained an equal volumetric proportion of unique WT (50%) and TET2 (50%) clones mixed together for competition against one another. Competition assays were cultured for 14 further days, for a total experiment duration of 28 days.

***Flow cytometry profiling of expanded clones***

At the endpoint of each culture (28 days), clone composition was profiled by flow cytometry. Between 20-25% of each clone was collected and transferred to a V-bottom polypropylene plate (Greiner Bio-one), followed by a centrifugation step (300 x g, for 5 minutes at room temperature) to remove culture media. Cells were resuspended in antibody cocktail prepared in FACS buffer, containing the following markers: EPCR PE, CD45.1 PE-Cy7, CD45.2 AF700, Sca-1 BV421, c-Kit (CD117) APC-Cy7, CD11b BV605 and Gr-1 FITC. Antibody excess was rinsed off by centrifugation, and cell pellets were resuspended into 7AAD (1:1000) for discrimination of viable cells from non-viable cells. Samples were analysed on a Cytoflex LX 355 or 375 or Cytoflex S (Beckman Coulter) using the high-throughput plate reader mode. Standardisation across cytometres was achieved by using a single fluorescence intensity bead (peak 6, Spherotech) with the CytoFlex built-in standardisation function within the CytExpert Software (Beckman Coulter), which enabled auto-calibration of the detector gain settings from the user-defined target values.

***Single-cell RNA-sequencing (scRNA-seq)***

For scRNA-seq analysis, split culture fractions were harvested on day 14 of culture, and transferred into sterile, 1.5-mL LoBind tubes (Eppendorf). Cell suspension volume was brought up to a total of 1mL with sterile PBS, 1X and centrifuged at 500 x g for 5 minutes at 4^o^C. Fixation and barcoding were performed according to the Parse Biosciences single-cell pipeline manufacturer’s protocol. Briefly, cells were fixed with the Evercode^TM^ Cell Fixation v. 2 kit (Parse Biosciences). Libraries were prepared using the Evercode WT v.2 kit (Parse Biosciences). Samples were then sequenced using the Illumina NovaSeq 6000 platform.

***scRNA-seq data analysis***

FASTQ files from 8 sublibraries were demultiplexed using the splitpipe pipeline based on unique barcodes (v.0.7.6, Parse Biosciences). Demultiplexed FASTQ files were aligned to the mouse reference genome (GRCm39) by the Cell Ranger method. Cell-specific barcodes were determined by the EmptyDrops method, integrated in the CellRanger pipeline (Lun et al, 2019); background-associated barcodes were excluded from analysis. Quality control and subsequent data analysis were performed using the Scanpy (Wolf et al, 2018) pipeline in Python (v. 3.14.6). The established criteria for quality control were as follows: 1) Cells with less than 1000 detected genes; 2) Cells with over 10% of unique molecular identifier (UMI) counts for mitochondrial genes; 3) Multiplets (which were excluded by the Scrublet pipeline (Wolock et al, 2019). Cells which passed these criteria were included in the next phase of analysis. Non-linear dimensionality reduction was performed by the Uniform Manifold Approximation and Projection (UMAP) embedding method, and executed using the Scanpy function “sc.pl.umap”. The Leiden algorithm (Traag et al, 2019) was applied using the “sc.tl.leiden” function of Scanpy for cell clustering.

***Bulk RNA-sequencing***

The control wells of the selected competitor clones were subjected to transcriptomic profiling, by harvesting 50% of clone contents at the experimental endpoint (28 days). RNA was extracted from clones using the PicoPure RNA kit (Applied Biosystems) by resuspending the samples into sterile PBS, 1X, centrifuging and incubating with RNA extracting reagent for 30 min at 56C on a thermoblock. The mixture was centrifuged; the supernatant was recovered, transferred to new collection tubes and stored at -80C. Samples were then resuspended in molecular biology-grade 70% Ethanol, transferred to the pre-conditioned columns and incubated for 2 min to enable RNA binding. Columns were washed up, and RNA was treated with DNAse-I (Qiagen) to ensure the purity and quality of the sample. Columns were washed in two more subsequent steps; Elution buffer was added to the columns, RNA was recovered and stored at -80^o^C. Quality of RNA samples was assessed on the Bioanalyzer 2100 (Agilent Technologies) by the Genomics Lab from the Technology Facility, University of York. Samples with poor quality or lower-than-threshold concentration were discarded and removed from the workflow. Libraries were prepared using the NEBnext Multiplex Oligos library prep kit and Unique Dual Index Primer Pairs for Illumina next generation sequencing (NGS) (New England Biolabs).

***Bulk RNA-seq data analysis***

Data were retrieved from Novogene or Azenta, and aligned to the mouse reference genome (GRCm39) by STAR method. Quality control of data was done by the CutAdapt trimming method v 5 (Martin, 2011). Gene counts were obtained through the Python Library HTSeq (High-Throughput sequencing) (Anders et al, 2015). Batch effect correction was performed on the data by performing a principal component analysis on samples obtained across different independent experiments, and no batch effect was observed. Prior to comparing across groups, log-transformed counts for each sample were obtained for the 100 most highly expressed genes across individual samples. Differential gene expression analysis was performed on RStudio by the DESeq 2 method (Love et al, 2014); a PCA was performed on all sample conditions. Estimation of log-fold change in gene expression was done by empirical Bayesian data shrinkage. Volcano plots of differentially expressed genes were produced using the ggplot2 and AnnotationDbi packages (CRAN repository). Kyoto Encyclopedia of Gene and Genomes (KEGG) pathway enrichment analysis was performed using the clusterprofiler package, and pathways were directly visualised on the KEGG website (https://www.genome.jp/kegg/pathway.html) using the pathview package (CRAN repository). Representative heatmaps of log2-transformed counts from ordered DESeq2 results were produced using the pheatmap package; single gene expression whisker plots were produced using the ggplot2 and ggpubr packages. Gene expression data was normalised using the median of ratios method in DESeq2 (Love et al, 2014). Overrepresentation analysis of Gene Ontology (GO) terms from differentially expressed genes was performed using clusterProfiler; data were visualised with the enrichplot package.

***Data-Independent Acquisition Mass Spectrometry (DIA-MS)***

Supernatants were collected from competition cultures on days 26 and 28 of the competition cultures, and transferred to polystyrene V-bottom plates. Supernatants were centrifuged for 5 min at 500 x g at 4°C, then carefully collected and transferred into new polypropylene V-bottom plates. Plates were stored at -80C until further use. Supernatants from two timepoints were pooled and transferred to 2-mL protein LoBind tubes (Eppendorf). Samples were precipitated with Ice-cold acetone at a 4:1 ratio, and were dissolved in ammonium bicarbonate (AmBic) (Sigma Aldrich). Disulfide bonds were reduced by incubation with dithiothreitol (DTT) (Sigma Aldrich) at 56^O^C for 30 min on a dry block heater. The alkylating agent iodoacetamide (IAA) (Sigma Aldrich) was added to each sample, which were incubated for 30 min at room temperature protected from light. 0.5 ug trypsin (Promega) was added to each sample and digestion was carried out at 37^o^C overnight. Digestion was stopped by addition of 10% TFA and samples were dried by vacuum centrifugation. Sample loading into EvoTips (EvoSep biosystems) was performed according to the manufacturer’s protocol. Samples were then introduced into the acquisition instrument on a 15-cm performance column (15 cm x 150 μm x 1.5 μm) with elution, using a pre-set 30 SPD standard method with a 44-min gradient. Data were acquired using the DIA-MS method on a Bruker timsTOF HT mass spectrometer, between 400-1200 mass-to-charge (m/z) ratio, using 25 Th DIA windows for each run. The nanoUPLC system was interfaced to a timsTOF HT mass spectrometer (Bruker) with a CaptiveSpray ionisation source. Positive PASEF-DIA, nanoESI-MS and MS2 spectra were acquired using CompassHyStar software (version 6.2, Bruker). Instrument source settings were capillary voltage, 1600 V; dry gas, 3 l/min; dry temperature; 180^o^C. Spectra were acquired between m/z 100-1700. DIA windows were set to 25 Th width between m/z 400-1201 and a TIMS range of 1/K0 0.6-1.60 V.s/cm2. Collision energy was interpolated between 20 eV at 0.65 V.s/cm^2^ to 59 eV at 1.6 V.s/cm^2^.

***DIA-MS Data analysis***

LC-MS data, in bruker .d format, was processed using Spectronaut (version 19) software and searched in direct-DIA mode against the mouse subset of Swiss prot appended with common proteomic contaminants. Search criteria specified: enzymes, trypsin/P; variable modifications; acetyl (protein N-term), oxidation (M); fixed modification, carbamidomethyl; digest type, specific; maximum peptide length, 52; minimum peptide length,7; missed cleavages, 2; MZ extraction strategy, maximum intensity; precursor Q value cutoff, 0.01. DIA XIC extraction windows for IM, IT, MS1 and MS2 were all set as dynamic. Identifications were filtered to require protein q-values < 0. 01 and a minimum of two peptides per accepted protein. An average of 5500 protein groups were reported across all samples. Normalisation of data was performed on R Studio using the trimmed means of M-values method (Robinson & Oshlack, 2010). Z-scores were calculated and data were log2-transformed, and representative volcano plots were made for significant differences in protein intensities using the ggplot and EnhancedVolcano packages. Secreted proteins (classified as peripheral, transmembrane, lipid anchor and soluble) were identified amongst the differentially expressed protein dataset by the DeepLoc protein localisation method as previously described (Ødum et al, 2024).

***HSC Transplantation***

From split cultures, 50% of total clone contents were collected, and from competition cultures, 30% of the clone contents were collected and prepared at the experimental endpoints (28 days for both experimental settings). Cells were transferred to 1.5-mL lo bind tubes (Eppendorf), topped up with PBS, 1X and centrifuged for 5.5 min at 300 x g at 4°C. Supernatant was carefully removed and samples were resuspended into 300 uL of sterile PBS, 1X for injections. For split cultures, Ly5-congenic CD45.1 (W41/W41) recipients were sub-lethally irradiated and reconstituted immediately after with the samples. For competition assays, CD45.1 or CD45.2 BL6 recipients were lethally irradiated, and reconstituted with competitor clones along with 2.5 x 105 whole bone marrow helper cells from hybrid CD45.1/CD45.2 donors. WT competitor donor cells (CD45.1) were transplanted into CD45.2 recipients, while TET2 competitor and TET2-super competitor donor cells (CD45.2) were transplanted into CD45.1 recipients. Monitoring of chimerism was performed by serial bleeding of recipients every four weeks up to week 24 post-injections. Peripheral blood (PB) samples were stained with CD45.1 PE-Cy7, CD45.2 AF700, CD3e PE, B220/CD45R APC, Ly6G BV421 and CD11b BV605 and were analysed on a BD Fortessa LSR X20. Hematopoietic tissues (bone marrow and spleen) were harvested from recipient mice at the end of the transplantation experiment, and analysed by flow cytometry.

***In vitro validation of targets***

Twenty-five ESLAM-Sca-1^high^ cells were sorted into individual wells in HSC expansion conditions supplemented with recombinant IFN-γ (MACS Miltenyi #130-105-785), Rho inhibitor (Rhosin) hydrochloride (Biotechne #5003/10), CDC42 activity inhibitor (CASIN) (Biotechne #5050/10) or NSC-23766 (Biotechne #2161/50) at the following concentrations: 2, 20, and 200 ng/mL IFN-γ; 2, 10, and 20 μM for CASIN; 1, 10, and 50 μM Rhosin; 5, 50 and 200 μM NSC-23766. Media was first changed on day 5, and then three times per week. Cells were analysed by flow cytometry at the 28-day endpoint. Data were exported as FCS files from a Cytoflex cytometre, imported into Flowjo for analysis, and processed in GraphPad Prism for statistical testing.
